# Diverse toxin repertoire but limited metabolic capacities inferred from the draft genome assemblies of three *Spiroplasma* (Citri clade) strains associated with *Drosophila*

**DOI:** 10.1101/2024.09.23.613922

**Authors:** Paulino Ramirez, Humberto Martinez Montoya, Rodolfo Aramayo, Mariana Mateos

## Abstract

*Spiroplasma* (Class Mollicutes) is a diverse wall-less bacterial genus whose members are strictly dependent on eukaryotic hosts (mostly arthropods and plants), with which they engage in pathogenic to mutualistic interactions. *Spiroplasma* are generally fastidious to culture *in vitro*, especially those that are vertically transmitted by their hosts, which include flies in the genus *Drosophila*. *Drosophila* has been invaded by at least three independent clades of *Spiroplasma*: Poulsonii (the best studied; contains reproductive manipulators and defensive mutualists associated two major clades of *Drosophila*; and has among the highest substitution rates within bacteria); Citri (restricted to the *repleta* group of *Drosophila*); and Ixodetis. We report the first genome drafts of *Drosophila*-associated Citri Clade *Spiroplasma*: strain *s*Moj from *D. mojavensis*; strain *s*Ald-Tx from *D. aldrichi* from Texas (newly discovered; also associated with *D. mulleri*); and strain *s*Hy2 from *D. hydei* (the only *Drosophila* species known to naturally also harbor a Poulsonii clade strain, thereby providing an arena for horizontal gene transfer). Compared to their Poulsonii clade counterparts, we infer that the three Citri clade strains have: (1) equal or worse DNA repair abilities; (b) more limited metabolic capacities, which may underlie their comparatively lower titers and transmission efficiency; and (c) similar content of toxin domains, including at least one ribosome inactivating protein (RIP), which are implicated in the Poulsonii-conferred defense against natural enemies. As a byproduct of our phylogenomic analyses and exhaustive search for certain toxin domains in public databases, we document the toxin repertoire in close relatives of *Drosophila*-associated *Spiroplasma*, and in a very divergent newly discovered lineage (i.e., “clade X”). Phylogenies of toxin-encoding genes or domains imply substantial exchanges between closely and distantly related strains. Surprisingly, despite encoding several toxin genes and achieving relatively high prevalences in certain natural populations (*s*Ald-Tx in this study; *s*Moj in prior work), fitness assays of *s*Moj (this study) and *s*Ald-Tx (prior work) in the context of wasp parasitism fail to detect a beneficial effect to their hosts. Thus, how Citri clade strains persist in their *Drosophila* host populations remains elusive.

**Data summary:** All novel sequencing data are available through National Center for Biotechnology Information (NCBI) repositories. Illumina raw reads, assemblies, and NCBI annotations are available under BioProject Nos. PRJNA506493 for sHy2, PRJNA506491 for sAld-Tx, and PRJNA355307 for sMoj. Oxford Nanopore (MinIon) reads for sHy2 are under SRA Accession Number SRR12348752.

Supporting Material is available under the DOI 10.6084/m9.figshare.c.7437997 or as accompanying supporting documents in the corresponding preprint server or scientific journal.

**Impact statement:** Symbiotic associations between arthropods and inherited microbes are pervasive, taxonomically and mechanistically diverse, and strongly influential. Research into the mechanisms and processes governing such heritable interactions is hindered by our inability to culture most inherited symbionts outside of their hosts. We studied three heritable strains of *Spiroplasma* (Citri clade) that naturally associate with *Drosophila* flies, and that reach relatively high prevalence in certain host populations, but appear to lack traits that would enable them to persist in host populations (e.g. such as high vertical transmission efficiency, reproductive manipulation, or fitness benefits). We compared their genomes to those of a separate *Spiroplasma* clade (Poulsonii) that associates with *Drosophila*, which does exhibit some of the traits that contribute to persistence, including protection against natural enemies of their hosts, and also has among the highest DNA substitution rates recorded for bacteria. Compared to Poulsonii, the three Citri clade strains have smaller genomes and fewer genes, leading us to predict they have similarly high DNA substitution rates, but more limited metabolic capacities, which may explain the comparatively lower densities they achieve within individual hosts, and their frequent loss in lab colonies of their hosts. However, the toxin repertoire of Citri clade was comparatively diverse, and the result of horizontal gene exchange among close and distant strains, and within-genome shuffling. We hypothesize that Citri clade strains persist via unknown fitness benefits conferred to their hosts, possibly mediated by toxins, or by substantial horizontal transmission. Our results, which also capitalized on publicly available assemblies, expand the range of *Spiroplasma* lineages that encode a particular combination of toxin types, and revealed the existence of a highly divergent lineage of *Spiroplasma* that associates with insects.

## 1 Introduction

Heritable associations between insects and bacteria are pervasive, influential, and taxonomically and functionally diverse[1]. Whereas a small proportion of insects engage in obligate (i.e., absolutely necessary for the host) mutualisms with heritable bacteria, over half of insect species are estimated to harbor facultative (i.e., non-essential) bacteria, whose predominant mode of transmission is via maternal transfer[1–3]. The impact of these heritable facultative bacteria, the most common of which is the genus *Wolbachia* (Alphaproteobacteria; intracellular), is far reaching and ranges from detrimental to beneficial phenotypes, including manipulation of host reproduction[male killing, cytoplasmic incompatibility, parthenogenesis induction, and feminization; 4], as well as increased and reduced susceptibility to environmental stressors and natural enemies[reviewed by 5]. Some of these phenotypes are being exploited for solving major human challenges, such as the use of *Wolbachia* to reduce dengue virus transmission by *Aedes* mosquitoes[6]. Because most heritable bacteria are practically impossible to culture outside the host[for some exceptions, see 7], comparative and functional genomics tools have proven invaluable for uncovering the mechanistic basis and evolutionary history of such phenotypes, particularly when hypothesized mechanisms can be further queried with the genetic tools afforded by model insects[e.g. the evolution and mechanistic basis of cytoplasmic incompatibility in Wolbachia; 8, 9-11]. Substantial research progress has also been achieved with other influential heritable bacteria of insects, including the Gammaproteobacterium *Hamiltonella* of aphids[e.g. 12, 13, 14], and members of the Class Mollicutes genus *Spiroplasma*.

*Spiroplasma* are small cell-wall-less bacteria strictly dependent on eukaryotic hosts, commonly arthropods and plants, with which they form pathogenic (e.g. the insect-vectored plant pathogens *S. citri* and *S. kunkelii*), commensalistic or mutualistic associations[15]. *Spiroplasma* are estimated to infect up to 7% of terrestrial arthropod species[16]. *Spiroplasma* inhabit both intra- and extra-cellular (e.g. insect hemolymph) environments. Under certain conditions/environments, many are motile and acquire the helical (spiral) shape implied by their name[15]. It was these features that revealed their presence in drops of *Drosophila* hemolymph observed under the microscope to early researchers, who initially thought they were spirochaetes[17]. The genus is composed of three large clades: Citri+Poulsonii+Chrysopicola+Mirum; Ixodetis; and Apis[18]. The Apis clade appears to lack vertically transmitted members, whereas the other two clades contain both vertically and horizontally transmitted members[19 and references therein]. Like other vertically transmitted endosymbionts, heritable *Spiroplasma* are fastidious[7, 20, 21]. While some progress has been made towards *in vitro* culture and transformation[22, 23], heritable *Spiroplasma* are not genetically tractable in practical terms. Nonetheless, the common association of heritable *Spiroplasma* with tractable insect hosts (e.g. several members of *Drosophila*, aphids, coccinelid beetles) has contributed to their establishment as a valuable system for the study of insect-bacteria interactions[24, 25].

Approximately 20 species of *Drosophila*, representing divergent clades, are known to naturally host *Spiroplasma*[26, 27, reviewed by 28]. Members of two three major *Spiroplasma* clades associate with *Drosophila*. Transgenerational transmission, interpreted as evidence of vertical transmission, has been demonstrated for *Drosophila*-*Spiroplasma* associations that have been assessed[reviewed by 19, 28]. Transovarial transmission, requiring *Spiroplasma* invasion of developing eggs from the hemolymph, has been demonstrated in the strain that naturally associates with the model organism *D. melanogaster*[29]. However, lack of congruence between host and symbiont phylogenies reveals multiple instances of horizontal acquisition of *Spiroplasma* by *Drosophila*[30, 31], and interspecific horizontal transmission via ectoparasitic mites, which are commonly found on wild *Drosophila*, has been demonstrated in the lab[32]. The fitness consequences of *Spiroplasma* infection for *Drosophila* are diverse: (a) neutral (or unknown); (b) beneficial in certain contexts, such as enhanced tolerance or resistance against particular natural enemies that include endo-macroparasites (parasitic wasps and nematodes)[e.g. 33, 34], as well as against bacteria and fungi[35]; (c) reproductive manipulation in the form of male killing[reviewed in 25]; and (d) detrimental in the form of shortened life span [36] or increased susceptibility to certain pathogens[37].

Most knowledge on the *Drosophila*-*Spiroplasma* association has been gleaned from the Poulsonii clade, which: (a) contains male-killing (referred to as “SRO” = Sex Ratio Organisms; in early studies) and non-male-killing strains of *Drosophila*; (b) achieves relatively high densities within individual hosts[38]; (c) exhibits relatively high vertical transmission fidelities; and (d) includes all of the known defensive *Spiroplasma* of *Drosophila[*with the exception of an unpublished report from the Ixodetis clade associated with Drosophila atripex; 19]. While male-killing strains of the Poulsonii clade tend to achieve low prevalence in wild populations, non-male-killing strains can achieve high prevalence[reviewed by 28], likely due to their defensive phenotype[e.g. 33]. Substitution rates in the Poulsonii clade are among the highest reported for any bacteria[39], a feature that likely explains the repeated loss of the male-killing phenotype in lab populations. Poulsonii also contains the only successfully *in vitro* cultured *Drosophila*-associated *Spiroplasma*, which facilitated the initial comparative/functional genomics and proteomics studies[23, 40–42]. With the aid of the heterologous expression tools provided by *D. melanogaster*, and to a lesser extent by *E. coli*[43], much has been elucidated about the host and *Spiroplasma* factors and mechanisms involved in the male-killing and wasp/nematode killing mechanisms[44–51]. In both phenotypes, *Spiroplasma*-encoded toxins, including ribosome inactivating proteins (RIPs; similar to ricin and Shiga toxin), deubiquitinases (OTUs), and/or Ankyrin repeats, are implicated (see <u>Results and Discussion</u>).

The Poulsonii clade is associated with members of the two major groups of *Drosophila* (i.e., subgenus *Sophophora*, which contains *D. melanogaster*, and subgenus *Drosophila*). In contrast, the much less studied *Drosophila*-associated Citri clade: (a) is restricted to members of *repleta*[27, 30, 31, 52], a subgenus *Drosophila* group that contains most cactophilic *Drosophila*[53, 54]; (b) exhibits comparatively lower titers[38]; (c) achieves lower vertical transmission fidelity, based on its frequent loss from lab cultures of its hosts; (d) is not known to kill males; and (e) can exhibit a range of infection frequencies in wild host populations[up to 85%; 52].

Based on sequences of the 16S rDNA gene, three distinct *Drosophila*-associated Citri clade strains were previously known[30, 31]: (1) *s*Moj in *D. mojavensis*; (2) *s*Ald-West (originally *s*Ald) in *D. aldrichi* from Tucson, Arizona and in *D. wheeleri* from Catalina Island, California; and (3) *s*Hy2 (= *s*Hyd2) in *D. hydei*. *D. hydei* is also host to the Poulsonii clade strain *s*Hy1 (= *s*Hyd, *s*Hyd, or *s*Hyd1 in other studies), a strain that is geographically widespread, including Great Britain and Japan[30, 31, 52, 55–58], and whose genome was recently reported[39]. Both *s*Hy1 and *s*Hy2 co-occur in several localities in the American continent (Arizona, and the Mexican states of Oaxaca, Sonora, Estado de Mexico), which is the native range of *D. hydei*[54], but have not been recorded within the same individual fly[27, 30, 31, 52]. Co-occurrence within the same individual host would provide an arena for gene exchange between the two *Spiroplasma* strains.

Here we report the discovery and infection prevalence of *s*Ald-Tx (a fourth *Drosophila*-associated Citri clade strain), detected in two sympatric (non-sister) species of cactophilic *Drosophila* (*D. mulleri* and *D. aldrichi*) in the Texas Hill Country region. We also report on the fitness consequences of its close relative *s*Moj (native to *D. mojavensis*) in the context of wasp parasitism. We use short reads (Illumina) from whole genome shotgun DNA libraries to assemble, annotate, and compare the draft genomes of sAld-Tx, *s*Moj, and their close relative *s*Hy2 (native to *D. hydei*). We perform phylogenomic analyses of available representatives of the Citri and Poulsonii clades, along with outgroup taxa. We compare the inferred metabolic and DNA repair capacities between the Citri and Poulsonii clade *Drosophila*-associated strains, and their close relatives. Finally, we analyze putative toxin genes in our target strains, as well as in recently released genome assemblies of numerous *Spiroplasma* strains (most reflect byproducts of genome projects aimed at their insect hosts), including what appears to be a new clade of *Spiroplasma* associated with insects (termed “clade X” pending a more formal assessment).

## 2 Methods

### 2.1 *Spiroplasma* taxon naming convention

Because most of the *Spiroplasma* species or strains referred to in this study have not been formally described, we adopt the common practice, which has also been used for *Wolbachia*, of referring to unnamed *Spiroplasma* strains/species by lower case “*s*” (for *Spiroplasma*) followed by the first few letters of their host’s species name (e.g “*s*Mel” for the *Spiroplasma* strain of *Drosophila melanogaster*). In some cases, we add a number or region identifier to such labels.

### 2.2 Specimen Sources

We used banana+yeast baits or sweep-netting over a compost to collect wild *Drosophila* and parasitic wasps at several locations in the Austin, San Marcos, and San Antonio areas of Texas (for *D. aldrichi* and *D. mulleri*; distinguished on the basis coloration pattern of the tergites; https://flybase.org/reports/FBim0000512.html and https://flybase.org/reports/FBim0000511.html’; \Patterson, 1943 #5324), at Catalina Island, California (for *D. mojavensis*), and in central Mexico (for *D. hydei*). Several of the wild-caught females were used to establish isofemale lines; one per each of *D. aldrichi*, *D. mojavensis*, and *D. hydei* were used to sequence the genomes of their naturally-occurring Citri clade *Spiroplasma* strains (<u>Supporting Table S1</u>). All insects were maintained on Banana-Opuntia food (<u>Supporting Protocol S3</u>) at 25°C (12:12 light:dark cycle).

To determine whether flies were infected with *Spiroplasma*, DNA extractions of individual whole files were subjected to PCR with the *Spiroplasma*-specific primers 16STF1 and 16STR1 [30] with annealing settings of touch-down 65–55°C, which target a ∼1368bp region of the 16S rRNA gene. A subset of *Spiroplasma*-positive samples was subjected to Sanger sequencing. To estimate prevalence of *Spiroplasma* in time and space (in *D. aldrichi* and *D. mulleri* from Texas), we counted the number of positive and negative individuals. DNA extractions that did not produce amplicons with the *Spiroplasma*-specific PCR, were subjected to PCR of the host-specific mitochondrial gene Cytochrome Oxidase I (COI) with primers HCO2198 and LCO1490[59]. Extractions that yielded no COI amplicon were deemed of inadequate DNA quality, and thus excluded. All PCRs included positive (template from known *Spiroplasma*-infected fly) and negative (no DNA template) controls. Because *D. hydei* is the natural host of a Citri clade (*s*Hy2) and a Poulsonii clade (*s*Hy1) *Spiroplasma*[31], positive 16STF1-16STR1 amplicons of *D. hydei* individuals were subjected to separate restriction digestion reactions, each containing a different enzyme that targets diagnostic positions between the two strains (<u>Supporting Table S2</u>).

Three DNA isolation methods were used. For *Spiroplasma* screening purposes, we used “squish prep” [60]to extract DNA from individual flies. For whole genome shotgun (WGS) Illumina sequencing, 1–2 grams of whole adult flies (approximately 1–2 weeks old) from *Spiroplasma*-positive *D. hydei* and *D. aldrichi* (<u>Supporting Table S1</u>) were collected for separate CTAB-Phenol based DNA extractions (<u>Supporting Protocol S1</u>). DNA from *s*Moj-infected *Drosophila mojavensis* was isolated with a chloroform-ethanol extraction protocol (<u>Supporting Protocol S2</u>) from hemolymph obtained by piercing the mesothoracic segment of approximately 300 individuals belonging to the infected isoline CI-33-15. Immediately after the piercing, ∼35-40 flies were placed into 0.5 ml microcentrifuge tubes previously pierced in the bottom, which was placed within a 1.5 ml microcentrifuge tube containing ∼20 ul phosphate buffered saline solution 1X (PBS buffer; 137 mM NaCl, 2.7 mM KCl, 10 mM Na2HPO4, 1.8 mM KH2PO4), and centrifuged at 7000 rpm (g=4.5) for 10 sec.

### 2.3 Fitness assays to evaluate effect of *Spiroplasma s*Moj on *Drosophila mojavensis*, in the context of parasitism by two wasps

We used the following procedures to evaluate the effect of *Spiroplasma s*Moj of larva-to-adult survivorship of *D. mojavensis* in the context of wasp parasitism. We used *D. mojavensis* isoline CI-33-15 (<u>Supporting Table S1</u>) to establish *Spiroplasma*-infected and *Spiroplasma*-free sub-isolines (i.e., those that naturally lost the infection). Flies were allowed to oviposit on Opuntia-banana medium for 48 hours, and transferred to a new vial for a second oviposition; after which they were removed and subjected to individual DNA extraction (squish-prep) and *Spiroplasma*-specific PCR with 16STF1 and 16STR1 primers to verify infection status. For the *Spiroplasma-*infected treatment, only vials (replicates) in which all such females were *Spiroplasma*-positive were retained. Subsequently, 30 second-instar larvae were collected and transferred to a new vial (i.e., replicate), where they were subjected to one of the following wasp treatments: no wasp control; *L. heterotoma* Lh14; and *Asobara* sp. w35 (<u>Supporting Table S1</u>). Those subjected to wasps were exposed to adult female wasps in a 1:6 wasp:larvae ratio for 24h. Fifteen replicates per each of the six combined treatments (i.e., *Spiroplasma* x wasp) were obtained (see <u>Results and Discussion</u>). Values of initial larvae, puparia, eclosing adult flies, and eclosing adult wasps were recorded.

Graphing of results and statistical analyses were performed with the statistical software R, version 4.1.2 (R Development Core Team 2018). Fly and wasp survival measures were analyzed by fitting a generalized linear model with binomial distribution (or quasibinomial distribution when there was evidence of overdispersion). Each wasp treatment was analyzed separately. The significance of the independent variable *Spiroplasma* was assessed with Analysis of Deviance Table (Type II tests), as implemented in the ‘car’ package. Raw data (<u>Supporting Dataset S1</u>) and R command lines used (<u>Supporting Command Line S1</u>) are available in Figshare.

### 2.4 Preparation and Sequencing of DNA libraries

Illumina libraries of DNA extractions from *s*Moj-infected *D. mojavensis*, *s*Ald-Tx-infected *D. aldrichi*, and *s*Hy2-infected *D. hydei* were submitted to the Texas AgriLife Genomics and Bioinformatics Services Facility (College Station, TX) for library preparation (whole genome shotgun; Illumina) and sequencing. The *s*Moj library (paired-end 100bp) was sequenced on the HiSeq® 2500 Sequencing System (Illumina, Inc.), whereas the *s*Ald-Tx and *s*Hy2 libraries (paired-end 150bp) were sequenced on the NovaSeq-6000 system within an S2 flowcell (Illumina, Inc).

In an attempt to enhance the assembly of strain *s*Hy2, we used the following protocols to obtain long reads on the MinIon platform (Oxford Nanopore v2.1). To attempt enrichment of *Spiroplasma*, we collected hemolymph from *s*Hy2-infected *D. hydei* (H25 strain), and passed it through a 70µm filter. DNA was isolated from the filtered hemolymph using a chloroform-ethanol protocol (<u>Supporting Protocol S2)</u>. Because the amount of DNA was insufficient for the library preparation protocol, we combined it (“spiked it into”) a DNA sample from the tephritid fruit fly *Anastrepha striata* (confirmed to be *Spiroplasma*-free). The combined DNA was prepared for Oxford Nanopore sequencing with the Nanopore SQK-LSK109 sequencing kit following the manufacturer’s recommendations. Nanopore was run on a Spot-on flow cell Mk 1 R9 (FLO-MIN106), MinION Mk1B sequencer, utilizing the Nanopore sequencing software Minknow v2.1 (Oxford Nanopore). Basecalling was performed with Albacore v2.3.1 (Oxford Nanopore) and adapter sequences were removed from resulting reads with PoreChop (https://github.com/rrwick/Porechop).

### 2.5 Genome Assembly

Illumina raw data were inspected for quality with Fastqc[61]. Then Trimmomatic v.0.39 [62] was used to remove adapter sequences and filter low quality sequences using default parameters. To remove non-target (i.e., non-*Spiroplasma*) host reads, each library was aligned with Bowtie2 v.2.3.2 [63] to the closest *Drosophila* genome assembly available (lacking *Spiroplasma* infection): *D. hydei* ASM278046v1 for *D. hydei*; *D. mojavensis* dmoj_caf1 for *D. aldrichi* and *D. mojavensis*. Unmapped reads from the *D. hydei*, *D. aldrichi*, and *D. mojavensis* libraries were separately subjected assembly with SPades v.3.12[64]. Geneious Prime 2020.2.2 (Biomatters Ltd.), was used to obtain a single consensus *de novo* assembly per library (using default parameters).

#### 2.5.1 Final Assembly Quality Control

To reduce the inclusion of chimeric contigs and assembly artifacts in the final assemblies of each library, a series of quality control steps were applied to the metagenomic sequencing data. Assembled contigs with a length of 200bp or more were subjected to blastn [65] to the NCBI nt database (Oct 2018). Contigs with high homology (E-value = 1E^-50, PI>20) to *Spiroplasma* or closely related taxa were considered to belong to the target *Spiroplasma* strain, and retained. The reads were aligned to these selected contigs with Bowtie2 v.2.3.2 (default parameters), and the results were visualized and processed in Geneious. Contigs containing a region (excluding repeat regions located in the middle of the contig) coverage below a certain threshold (30x for *s*Hy2; 15x for *s*Ald-Tx and *s*Moj) were then broken apart at that site via the “Generate Consensus Sequence” option of Geneious. Nucleotide locations composed of more than 1 type of base were called as base if that base was in at least 50% of the reads mapped to that position. Otherwise, the consensus was assigned the corresponding IUPAC ambiguity code at that position. The resulting contigs are considered to be the final draft assembly for *s*Ald-Tx and *s*Moj.

The MinION trimmed fastq output (*s*Hy2) was assembled with the long read assembler CANU[66]. All Contigs were then subjected to a blastn search against the NCBI nt database (Oct 2018), resulting in 1 contig with a hit to *Spiroplasma*. The *s*Hy2 (short-read) SPAdes assembly was then aligned to the *Spiroplasma* CANU contig (Geneious Mapper), and aligned contigs were then merged with the CANU contig. *s*Hy2 Illumina reads were aligned to the remaining SPAdes contigs and the Canu-assembled contig with Geneious Mapper (5x iterations) with the purpose of correcting MinION-induced indels; a consensus was generated with the same settings as the *s*Hy2 SPAdes-only assembly. The resulting consensus contigs represent the final draft assembly for *s*Hy2.

### 2.6 Annotation

*s*Ald-Tx, *s*Hy2 and *s*Moj genome assemblies were submitted to the NCBI PGAP pipeline to obtain final annotations (Table 1). We used the following additional annotation tools. Protein sequences for the *Spiroplasma* genomes were annotated using the blastKoala KEGG tool v.2.2[September 2019; 67]. The generated KO numbers were then compared among several *Spiroplasma* genomes to determine differences in metabolism and DNA repair. Additionally, protein sequences were annotated for Cluster of Orthologous Groups (COG) by EGGNog v.5.0[68]. To assess completeness, the NCBI PGAP protein sequences were subjected to BUSCO v.5.5.0 (Benchmarking Universal Single-Copy Orthologs) analyses, as implemented in the usegalaxy.eu server (mode = prot; Lineage data source = Cached database with lineage all+2024-03-21-114020; lineage = Mollicutes and Entomoplasmatales). Genes with KOs of interest were compared via blastn to a nucleotide database composed exclusively of *Spiroplasma* strains (<u>Supporting Table S3</u>), to confirm their presence/absence in the Citri and Poulsonii clades.

**Table 1.**
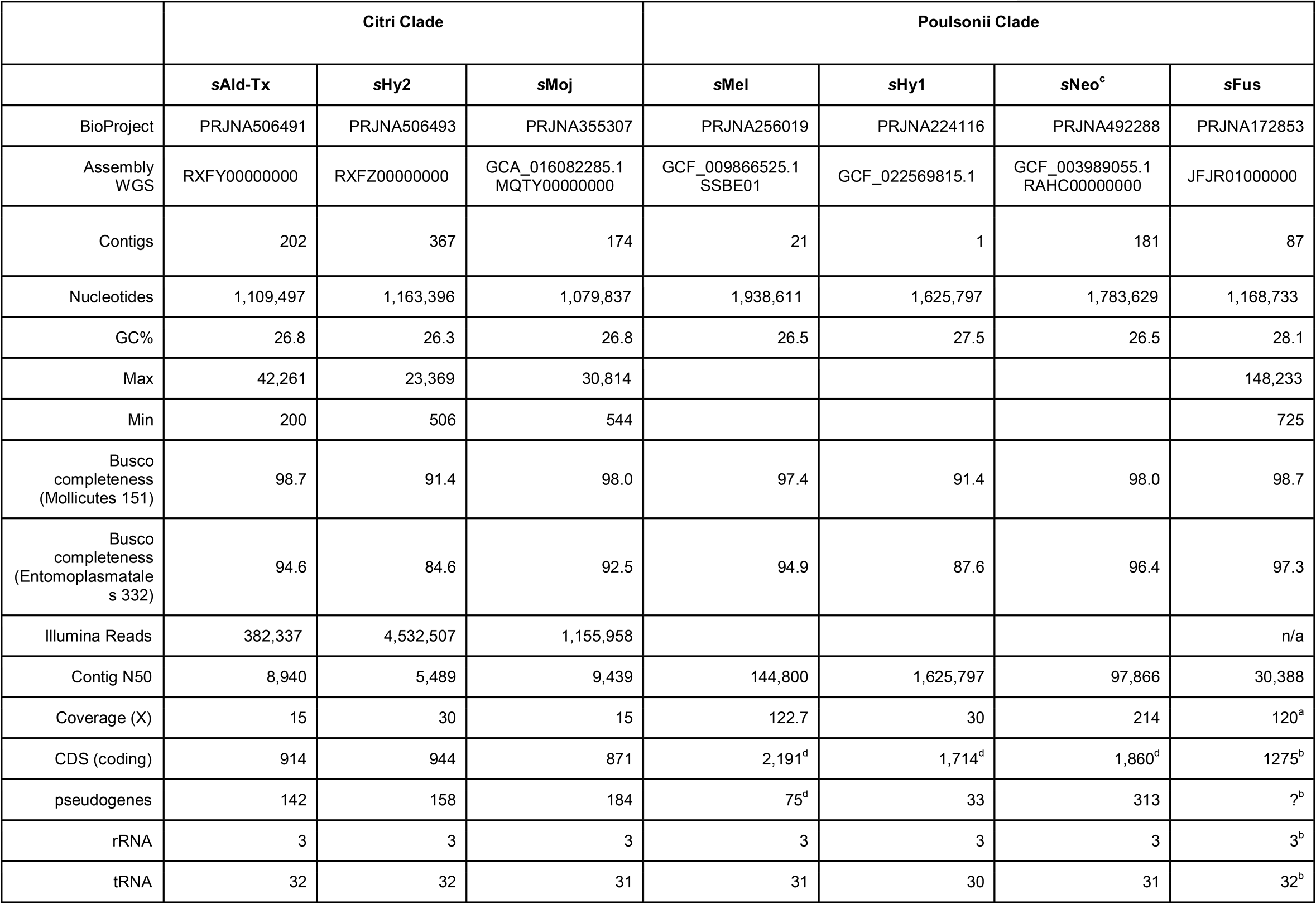
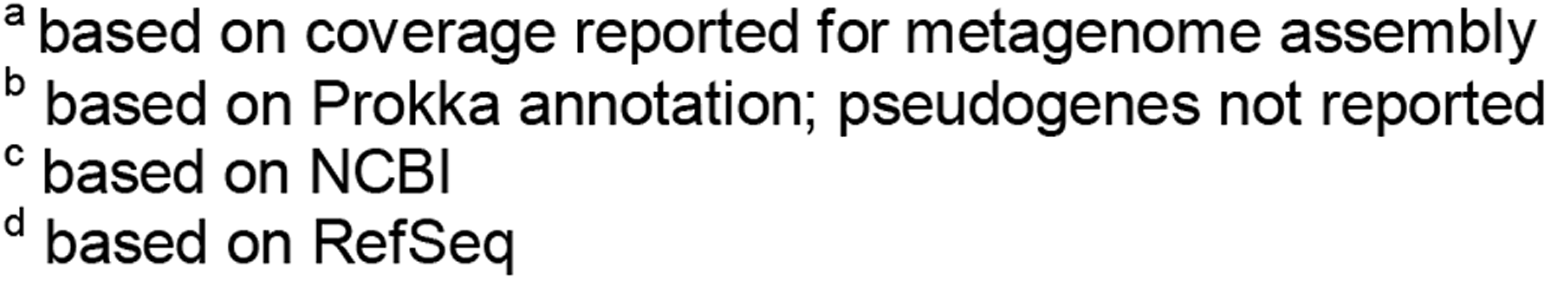
Assembly stats (excluding known plasmids) for the three *Drosophila*-associated Citri Clade strains, for *s*Fus (from *Glossina fuscipes*), and the previously reported *Drosophila*-associated Poulsonii Clade (for comparison).

|  | Citri Clade |  |  | Poulsonii Clade |  |  |  |
| --- | --- | --- | --- | --- | --- | --- | --- |
|  | sAld-Tx | sHy2 | sMoj | sMel | sHy1 | sNeo<sup>c</sup> | sFus |
| BioProject | PRJNA506491 | PRJNA506493 | PRJNA355307 | PRJNA256019 | PRJNA224116 | PRJNA492288 | PRJNA172853 |
| Assembly WGS | RXFY000000000 | RXFZ000000000 | GCA_016082285.1<br>MQTY000000000 | GCF_009866525.1<br>SSBE01 | GCF_022569815.1 | GCF_003989055.1<br>RAHC000000000 | JFJR010000000 |
| Contigs | 202 | 367 | 174 | 21 | 1 | 181 | 87 |
| Nucleotides | 1,109,497 | 1,163,396 | 1,079,837 | 1,938,611 | 1,625,797 | 1,783,629 | 1,168,733 |
| GC% | 26.8 | 26.3 | 26.8 | 26.5 | 27.5 | 26.5 | 28.1 |
| Max | 42,261 | 23,369 | 30,814 |  |  |  | 148,233 |
| Min | 200 | 506 | 544 |  |  |  | 725 |
| Busco completeness<br>(Mollicutes 151) | 98.7 | 91.4 | 98.0 | 97.4 | 91.4 | 98.0 | 98.7 |
| Busco completeness<br>(Entomoplasmatals 332) | 94.6 | 84.6 | 92.5 | 94.9 | 87.6 | 96.4 | 97.3 |
| Illumina Reads | 382,337 | 4,532,507 | 1,155,958 |  |  |  | n/a |
| Contig N50 | 8,940 | 5,489 | 9,439 | 144,800 | 1,625,797 | 97,866 | 30,388 |
| Coverage (X) | 15 | 30 | 15 | 122.7 | 30 | 214 | 120<sup>a</sup> |
| CDS (coding) | 914 | 944 | 871 | 2,191<sup>d</sup> | 1,714<sup>d</sup> | 1,860<sup>d</sup> | 1275<sup>b</sup> |
| pseudogenes | 142 | 158 | 184 | 75<sup>d</sup> | 33 | 313 | ?<sup>b</sup> |
| rRNA | 3 | 3 | 3 | 3 | 3 | 3 | 3<sup>b</sup> |
| tRNA | 32 | 32 | 31 | 31 | 30 | 31 | 32<sup>b</sup> |

<sup>a</sup> based on coverage reported for metagenome assembly <sup>b</sup> based on Prokka annotation; pseudogenes not reported <sup>c</sup> based on NCBI <sup>d</sup> based on RefSeq

In the process of annotating these genomes, we discovered and extracted 87 contigs (<u>Supporting Table S4</u>) in the draft genome assembly of the tsetse fly *Glossina fuscipes fuscipes* (PRJNA172853, WGS JFJR00000000.1) that we assigned to *Spiroplasma* on the basis of blastn (NCBI nt database Oct 2018). We also learned of the recent release of numerous *Spiroplasma* genome assemblies obtained from sequencing projects aimed at their arthropod hosts, many of which encode putative toxin genes (e.g. search for “spiroplasma [organism] AND inactivating” at https://www.ncbi.nlm.nih.gov/protein). The majority of these genomes had gene annotations available in NCBI. A preliminary phylogenetic analysis of the 16S ribosomal RNA gene allowed us to identify the major *Spiroplasma* clades to which they belong (<u>Supporting Figure S1</u>; <u>Supporting Dataset</u> <u>S2</u>). A small number of these genomes fell within the Citri and Poulsonii clades, and were thus included in the phylogenomic analyses. Because most of the remaining genomes fell within more distantly related clades (e.g. Apis, Ixodetis, and a previously unknown clade hereafter referred to Clade X), they were not included in the phylogenomics analyses, and were only partially examined for other features of interest (e.g. toxin genes). Establishing the phylogenetic position of Clade X is beyond the scope of this study, but based on <u>Supporting Figure S1</u>, it appears more closely related to Ixodetis than to Apis or Citri+Poulsonii+Chrysopicola+Mirum.

We used Prokka v.1.14 [69] with translation table 4 (Mycoplasmas) implemented within the Galaxy Project platform [70] to identify all putative protein coding regions in genome assemblies lacking publicly available gene annotations, and in all Citri and Poulsonii clades genomes used in the phylogenomics analyzes. Representatives of a sister lineage (the Chrysopicola clade), as well as the outgroup (Mirum clade), were also included (<u>Supporting Dataset S3</u>). Predicted protein sequences were analyzed with Interpro v. 5.25 [71, 72] to identify putative domains, implemented in usegalaxy.eu and https://www.ebi.ac.uk/interpro/ servers, using all the “Member databases” and “Other sequence features” available with their preconfigured cut off thresholds.

To search for genes encoding potential toxins, we searched the annotations (Interpro, Koala, and NCBI) for the following terms, most of which reflect toxins found in *Spiroplasma* or in other arthropod endosymbionts[reviewed in 73]: “toxin”; “lethal”; “inactivating” and “ricin” (for Ribosome Inactivating Protein); “etx”, “mtx”, “pore” (for certain pore-forming proteins), “ankyr” (for Ankyrin); “fic” (for filamentation induced by cAMP), “adp”, “protective” and “antigen” (for ADP-ribosyltransferase exoenzyme and its associated protective antigen), “cdt” (for cytolethal distending toxin); “cif” (for the cytoplasmic incompatibility factor); “pqq” (for PQQ-binding-like beta-propeller repeat protein) and “latro” (for latrotoxin). Genes with annotations associated with potential toxins, were subjected to further analysis to infer their evolutionary history and potential function. A subset of inferred protein products were determined to be truncated at the N or C termini, as they only contained single domains that are usually found within larger multi-domain proteins. To determine if a putative gene of interest was broken up by a missense mutation causing truncated single domain gene products, we searched for the “missing” regions/domains in the immediate flanking regions on their 5’ and 3’ ends. Similarly, because of the fractured nature of the assemblies in the Citri clade strains *s*Moj, *s*Ald-Tx, and *s*Hy2, and detection of intact and pseudogenized transposases in several contigs (see Results), we also searched for potentially missing domains of genes in other contigs (e.g., in cases where the end of a contig occurred within a predicted partial protein of interest).

Because a search for the term “inactivating” in the gene annotations revealed the presence of RIP genes of eight out of nine Clade X strains, and several of these genes shared substantial similarity with those of clades Citri and Poulsonii, we analyzed Clade X genome assemblies with Prokka and Interproscan to find all predicted proteins with domains of interest. For the Ixodetis and Apis clade, we extracted the genes of interest based on term search in their NCBI gene annotations only.

For certain genes of interest, such as toxins, we extracted specific domain regions from the amino acid sequences, aligned them with the MAFFT v.7.490 ([74, 75]plugin of Geneious, and performed phylogenetic analyses with IQTREE 2.2.2.6 COVID-edition for Mac OS X 64-bit built May 27 2023[76,Kalyaanamoorthy, 2017 #12197, 77]. For a subset of genes and taxa, we also performed phylogenetic analyses on the nucleotide sequences (see Results).

Strains for which the term “crispr” and/or “cas9” was detected in their annotation were analyzed with CRISPRCasTyper v1.8.0[78], as implemented in the webserver https://crisprcastyper.crispr.dk/; accessed 15 July 2024; default settings).

To further examine genes of potential plectroviral origin, we used WP_339038695.1 (product = plectrovirus svts2 rep protein) as a query for a Delta-blast search [79] against NCBI’s non-redundant database.

### 2.7 Phylogenomic Analyses

To identify single-copy orthologs, the Prokka-derived amino acid sequences of 35 genomes (<u>Supporting Dataset S3</u>) were analyzed with Orthofinder2 v.2.5.5 [80] (default parameters), and subsequently aligned with MAFFT v7.471 (-L-INS-I, with the - phylipout option). Gene loci alignments were assessed for recombination with PhiPack[81], utilizing windows of 10, 20, 30, 40, and 50 amino acids. Gene loci with a p-value of 0.05 or less with one or more of the window sizes were considered to have significant recombination and removed from further analysis. The remaining genes were concatenated and subjected to Maximum Likelihood phylogenetic analysis with IQ-TREE. Trees from these and the remaining analyses were visualized and edited in FigTree v1.4.4 (http://github.com/rambaut/figtree/). Trees and other figures were further edited in Inkscape (https://inkscape.org/).

## 3 Results and Discussion

### 3.1 Identity and infection frequencies of a newly discovered *Spiroplasma* strain (*s*Ald-Tx) in wild populations of *Drosophila aldrichi* and *Drosophila mulleri* from Texas

The 16S rDNA sequences of *Spiroplasma* from *D. aldrichi* and *D. mulleri* from Texas species were identical to each other, and were (1 out of 973; or 11 out of 1293 bp = 0.85% uncorrected p distance) different from the strain previously reported in *D. aldrichi* and *D. wheeleri* from California (*s*Ald-West; GenBank Acc. Nos. FJ657236 and FJ657227; <u>Supporting Figure S2</u>; <u>Supporting Dataset S4</u>). Hereafter, we refer to the *Spiroplasma* strain associated with *D. aldrichi* (and *D. mulleri*) from Texas as *s*Ald-Tx (or *s*Ald-East).

The infection prevalence of *s*Ald-Tx varied broadly (4% to 94%) across its two host species, sites and years, but infection prevalence was consistently higher in *D. aldrichi* compared to *D. mulleri* (overall 71 vs. 38%, respectively; <u>Supporting Table S5</u>). These infection frequencies are within the range reported for other *Drosophila*-associated Citri clade strains: 33 and 85% in *D. mojavensis* (Arizona and California, respectively); 5.5% in *D. aldrichi* from Arizona; and 53% in *D. wheeleri*. *s*Hy2 prevalence values in *D. hydei* are unknown, as most studies only detected *s*Hy1 [55–58, 82] or did not separate *s*Hy1 from *s*Hy2 when computing *Spiroplasma* frequencies[52]. The highest reported *Drosophila*-associated *Spiroplasma* prevalence is that of *s*Neo (Poulsonii clade) in certain populations of *D. neotestacea*, where substantial fitness benefits in the form of protection against nematodes and parasitoid wasps, and no reproductive manipulation has been reported[33, 83]. In contrast, the infection frequencies of *Drosophila*-associated male-killing *Spiroplasma* strains tend to be <5%[17, 84–86].

### 3.2 No evidence of fitness benefits from *Drosophila*-associated Citri clade *Spiroplasma*

Our results revealed that *s*Moj does not significantly affect the larva-to-adult survivorship of its native host *D. mojavensis* following exposure to one of two divergent parasitic wasps (Figure 1a, c, d; <u>Supporting Table S6</u>): the braconid Aw35; and the generalist figitid *Leptopilina heterotoma* (strain Lh14). Strain *s*Moj also did not significantly affect success of these wasps developing in *D. mojavensis* (Figure 1b; <u>Supporting Table S6</u>). In contrast, wasp Lh14 is highly susceptible to *Spiroplasma* from the Poulsonii clade in several different *Drosophila* species[34, 83, 87–90]. Wasp Aw35 has not been tested against Poulsonii clade strains. Similarly to *s*Moj, Martinez-Montoya [27]reported that *s*Ald-Tx in *D. aldrichi* does not offer protection against wasps Aw35 and Lh14. The effect of *s*Hy2 on *D. hydei*, and of *s*Ald-Tx in its other native host *D. mulleri*, in the context of wasp exposure has not been established. It is possible that these Citri clade strains protect against other natural enemies, particularly in light of their encoding of several putative toxin genes (see below). There is a huge known and predicted diversity of *Drosophila* parasitoids[91], of which only ∼10 species have been assessed for susceptibility to one or more *Spiroplasma* strains[27; and references therein, 92, 93]. Our single attempt to sample *Drosophila* parasitoids in Catalina Island, California, where *s*Moj achieves high prevalence, failed, but several *Drosophila* parasitoid species, including Aw35, have been sampled in the habitat of *D. aldrichi* and *D. mulleri* in Texas[91]. In addition, it is possible that the Citri clade strains studied herein protect against other types of natural enemies (e.g. bacteria, viruses, protozoans, fungi), as enhanced host fitness against pathogenic fungi and/or bacteria has been documented for other *Spiroplasma* clades[35, 94].

**Figure 1.**
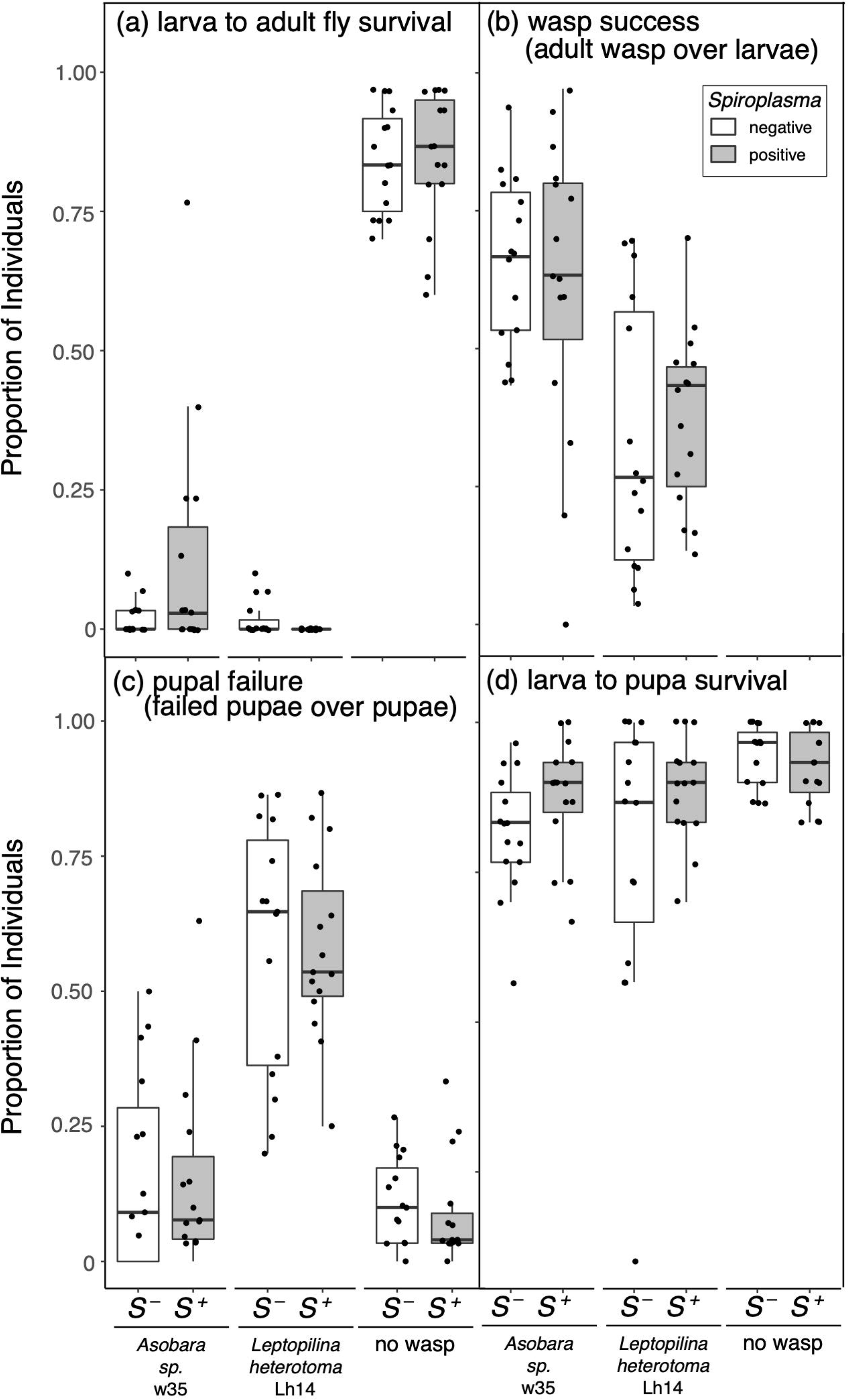
Effect of *Spiroplasma sMoj* (in *Drosophila mojavensis*) on selected fly and wasp success and failure measures: (a) larva to adult fly survival; (b) wasp success; (c) pupal failure; and (d) larva to pupa survival. Points represent each measurement obtained. Box plots display the median, upper and lower quartiles, and the range excluding points beyond 1.5 * IQR (inter-quartile range). Grey boxes = *Spiroplasma*-infected (*S*^+^); White boxes = *Spiroplasma*-free (*S*^−^). Wasp treatments (*Asobara* sp. w35, *Leptopilina heterotoma* Lh14, and no wasp control) are indicated in the X-axis.

In the absence of wasps, Citri clade *Spiroplasma* strains *s*Moj and *s*Ald-Tx have weak to no effect on larva-to-adult fly fitness[Figure 1; and 27]. Reproductive phenotypes have been investigated for *s*Moj and *s*Hy2[27], who ruled out Cytoplasmic Incompatibility (CI) in *s*Moj, and found that *s*Moj-infected *D. mojavensis* and *s*Hy2-infected *D. hydei* tend to lay more eggs earlier than their *Spiroplasma*-free counterparts. In contrast, the Poulsonii clade strain *s*Hy1 does not exert a detectable effect on *D. hydei* oviposition[95]. Increased early oviposition was reported for the Poulsonii clade (male-killing) strain (WSRO) harbored by *D. willistoni*[96]. An early mating propensity induced by the Poulsonii clade strain (NSRO) harbored by *D. nebulosa* was reported byMalogolowkin-Cohen and Rodrigues-Pereira [97]. No further fitness consequences of Citri clade strains on *Drosophila* have been examined. Unfortunately, experimentation with Citri clade strains has been more challenging than that with Poulsonii clade strains, due to the frequent unintentional loss within their native hosts, and the difficulties of artificially transferring and maintaining Citri clade strains over several generations[98; and personal observation]. Such difficulties might stem from the comparatively lower titers of the Citri clade (*s*Moj and *s*Hy2) vs. Poulsonii clade[sHy1 and the male-killer sMel; 38]. How *s*Ald-Tx and *s*Moj strains can achieve relatively high infection frequencies in wild populations despite low vertical transmission efficiency and no evidence of net fitness benefits, remains unknown, and could rely on substantial horizontal transfer.

### 3.3 Phylogenomic Relationships

We used phylogenomic analyses to infer the evolutionary history of currently available representatives of the Citri and Poulsonii clades plus outgroup taxa (<u>Supporting Dataset</u> <u>S3</u>). Three of the 35 genomes assemblies targeted for our phylogenomics inferences (i.e., strains *s*Ama, *s*Rhe, and *s*Cru) appeared substantially incomplete. Because their inclusion led to a small number (n = 41) of single-copy orthologues, we excluded them from the main phylogenomics dataset, which encompassed 32 taxa, and 189 genes lacking evidence of recombination. The concatenated alignments of the amino acid sequences of these 189 genes resulted in 62,325 positions, containing 33,279 distinct patterns, and 24,622 parsimony-informative sites. The inferred tree (Figure 2; <u>Supporting Dataset S5</u>) reveals that the *s*Ald-Tx, *s*Hy2, and *s*Moj strains form a monophyletic group, with *s*Moj as sister to *s*Ald-Tx + *s*Hy2. This relationship is consistent with inferences from a smaller number of genes[30, 31]. This relationship does not match the host phylogeny, because *D. mojavensis, D. mulleri* and *D. aldrichi* belong to the *mulleri* complex, which excludes *D. hydei*[99], implying horizontal transfer of *Spiroplasma* within this group of *Drosophila*. Additional evidence consistent with horizontal transmission includes the sharing of strain *s*Ald-Tx by sympatric specimens of *D. aldrichi* and *D. mulleri*; two species that are not sisters and are estimated to have diverged ∼5.56 mya[99]. Whereas the sharing of strain *s*Ald-West by *D. wheeleri* from California and *D. aldrichi* from Arizona [30, 52] could reflect another instance of horizontal transfer, taxonomic uncertainty includes the possibility that these two hosts are sister lineages whose common ancestor harbored *s*Ald-West[100–102]. Possible routes of horizontal transmission include via ingestion, and vectored by ectoparasitic mites or parasitoid wasps. There are documented examples of different cactophilic *Drosophila* species sharing the same species of mites[103], of *Drosophila* mites harboring *Spiroplasma*[104], as well as of interspecific transmission of *Spiroplasma* through mites[32]. Whereas no study has tested *Spiroplasma* transmission via parasitoid wasps, Gehrer and Vorburger [105]demonstrated this transmission mechanism for the hemolymph-dwelling symbionts *Hamiltonella defensa* and *Regiella insecticola* of aphids. One study reports horizontal transmission of *Spiroplasma* (Poulsonii clade) in *Drosophila* via ingestion, but attempts to repeat this finding failed[17, 96, 106].

**Figure 2.**
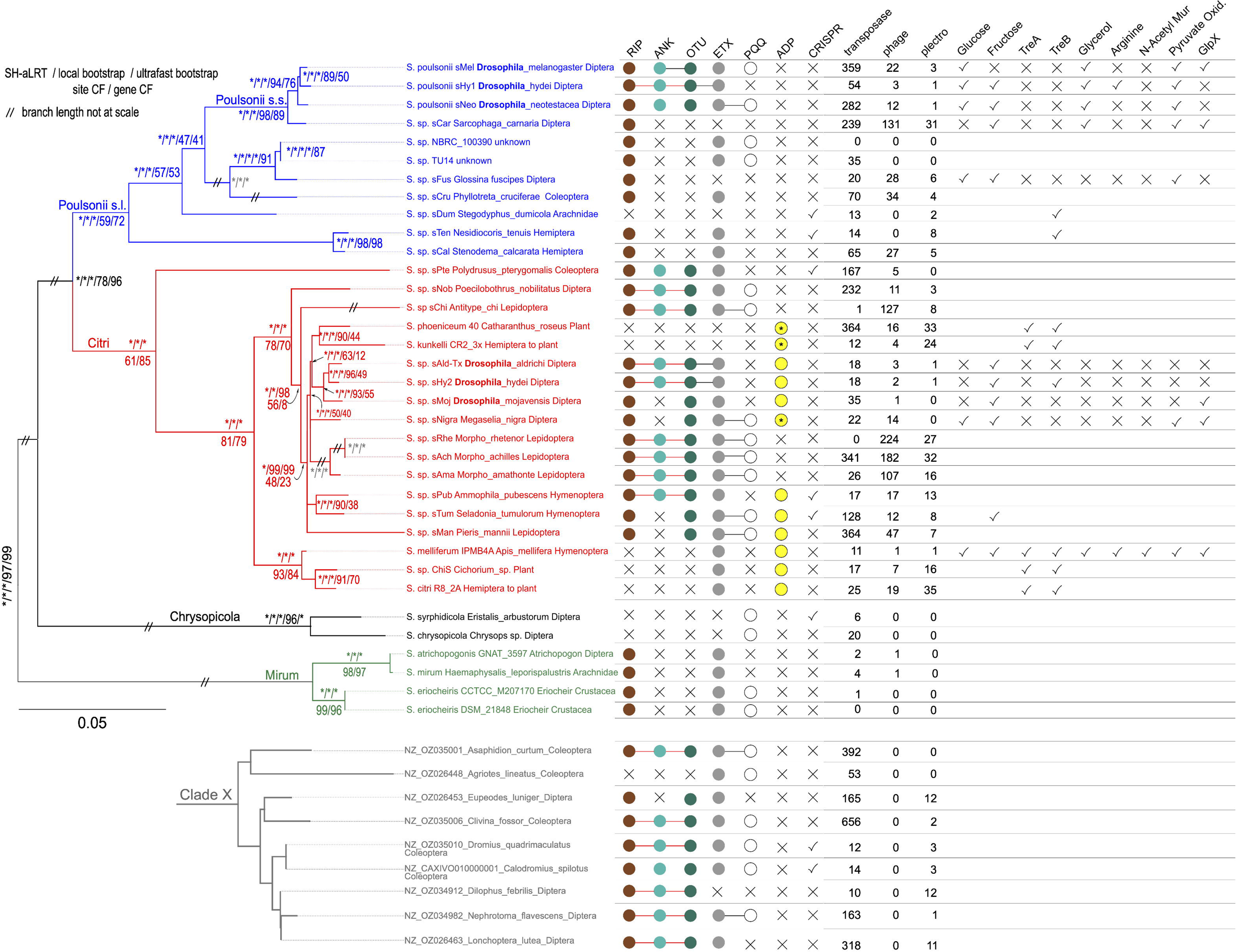
Phylogenomic relationships of members of the Citri and Poulsonii clades and outgroups, and 16S rDNA phylogeny of members of Clade X. Phylogenetic distribution of genes of interest. Tree is based on the 62,325 amino acid sites from 189 single-copy genes with no evidence of recombination, except for the position of sCru, sRhe, sAma (inferred based on a subset of 16,372 amino acid sites from 41 single-copy genes; first three support values in gray font). Clade support values are shown next to nodes (asterisks = 100% support). Tip labels contain the *Spiroplasma* species or strain name, followed by the host Genus_species, and its major taxonomic group. Clade X tree (gray) is based on NJ distance analysis of the 16S rDNA gene only. Circles to the right indicate presence of genes with domains annotated as RIP, Ankyrin (ANK), OTU, ETX, PQQ, and ADP (asterisks indicate that such gene is only found as pseudogenized). Circles connected by a single color line indicate that they occur within a single CDS at least once in the corresponding genome. An X indicates absence of such a gene. Columns with headers “transposase”, “phage”, “plectro” report the number of genes with such annotations. For the remaining columns, a check mark indicates presence, an X indicates absence, and a blank indicates its presence/absence was not determined.

The closest sister lineage to these *Drosophila*-associated Citri clade strains is the plant-pathogenic clade formed by *S. phoeniceum* and *S. kunkelli*. This *Drosophila*-associated plus plant pathogenic clade is joined at a polytomy with *s*Nigra[from the mushroom-feeding phorid fly Megaselia; 107], *s*Man[from the southern small white butterfly; 108], a clade of three *Morpho* butterfly associates[sAch, sRhe, and sAma; 109], and a clade associated with Hymenoptera (*s*Pub and *s*Tum). More distantly related strains associated with Coleoptera (*s*Pte), other Diptera (*s*Nob), other Lepidoptera (*s*Chi), and a clade formed by *S. citri*, *S. melliferum*, and ChiS, were recovered, and collectively assigned to the Citri clade (red). Sister to Citri is what we refer to as the Poulsonii *sensu lato* clade (blue). Poulsonii *s. l.* contains a subclade composed of the previously characterized *Drosophila*-associated strains (*s*Mel, *s*Hy1, and *s*Neo). Sister to this *Drosophila*-associated clade is *s*Car, associated with another dipteran, the flesh fly *Sarcophaga carnaria*. This Poulsonii *sensu stricto* clade is sister to a clade containing previously characterized (and *in vitro* cultivated) strains from unknown hosts NBRC_100390 and TU14 [110, 111] and two strains whose genomes have not been previously compared (*s*Fus from a tsetse fly and *s*Cru from a beetle). The most basal split in Poulsonii *s. l.* separates all of the above strains from two Hemiptera-associated strains (*s*Ten and *s*Cal).

### 3.4 Genome Assembly Statistics and Annotations

The assembly sizes of the three *Drosophila*-associated Citri clade strains (*s*Ald-Tx, *s*Moj, and *s*Hy2) ranged from 1.08 to 1.16Mb (Table 1), which is substantially smaller than those of their closely related plant-pathogens (*S. kunkelli*: 1.5Mb; *S. phoeniceum*: 2.2Mb), and of the *Drosophila*-associated Poulsonii clade strains (1.6–1.9Mb). The assemblies of *s*Ald-Tx, *s*Moj, and *s*Hy2 were substantially fragmented, ranging from 174 (*s*Moj) to 367 (*s*Hy2) contigs, and contig N50 was small (5.5 Kb in *s*Hy2 to 9.4 Kb in *s*Moj), compared to that of Poulsonii clade genomes (.98–1.6Mb), which did not solely rely on short reads. The fragmented status of these assemblies precludes conclusive inferences about genome sizes, synteny, and distribution and number of mobile/repetitive elements. The NCBI PGAP pipeline annotated 914, 944, and 871 protein-coding sequences for *s*Ald-Tx, *s*Hy2, and *s*Moj, respectively (Table 1), which are substantially fewer than those in their Poulsonii clade counterparts. Busco completeness scores using the Mollicutes and Entomoplasmatales lineage databases (containing 151 and 332 groups, respectively) were lowest in *s*Hy2 (91.4 and 84.6%) and similarly high in *s*Moj (98 and 92.5%) and *s*Ald-Tx (98.7 and 94.6%). These scores are similar to most other assembled *Spiroplasma* genomes[e.g. 107, 112, 113]. The number of tRNAs was similar among the assemblies (32 in *s*Ald-Tx and *s*Hy2; 31 in *s*Moj). Pseudogene content ranged from 142 (sAld-Tx) to 184 (*s*Moj), which is substantially larger than in *s*Mel (75) and in *s*Hy1 (33), but not in *s*Neo (313; see Table 1).

The assembly size of the Poulsonii *sensu lato* strain *s*Fus (from 87 contigs included with the *Glossina fuscipes fuscipes* assembly) was also relatively small (∼1.17 Mb; Table 1), but not much smaller that the complete chromosome size (∼1.2 Mb) of its sister lineage (comprised of strains TU-14 and NBRC_100390). Prokka annotated 1275 protein-coding genes in *s*Fus, compared to 1039 in TU-14 and NBRC_100390, but the number of pseudogenes in *s*Fus has not been determined. Busco completeness scores for *s*Fus using the Mollicutes and Entomoplasmatales lineage databases were relatively high (98.7 and 97.3%, respectively).

#### 3.4.1 Extrachromosomal and mobile genetic elements, and CRISPR/Cas system

Mobile genetic elements such as plasmids, phages, and insertion sequence (IS) elements, are common in numerous *Spiroplasma* genomes and other insect symbionts, and contribute to acquisition (and loss) of relevant functions (e.g. virulence factors) and rapid genome evolution[14, 39, 73]. While a full picture of such elements requires complete assemblies, which are lacking for the three *Drosophila*-associated Citri clade strains, below we discuss preliminary inferences. It is unclear whether the three *Drosophila*-associated Citri clade strains contain plasmids, as our assemblies failed to recover circular contigs. However, the three assemblies contained contigs encoding a *ParA family protein*: one in *s*Moj; and six in *s*Hy2 and *s*Ald-Tx; with one of each contig per strain also containing a *ParB* gene immediately downstream of the *ParA* gene. ParA and ParB are commonly involved in plasmid segregation[114]. Some of these contigs exhibit partial blast matches to known Citri and Poulsonii clade plasmids, but they also match parts of chromosomes (not shown). Concerning genes annotated as transposases, a substantially smaller number was detected in the *Drosophila*-associated Citri clade (*s*Ald-Tx = 18; *s*Hy2 = 18; *s*Moj =35) than in its Poulsonii clade counterpart (*s*Mel = 359; *s*Neo = 282; *s*Hy1= 54) (Figure 2), but complete assemblies may reveal more.

Previous studies indicate that the Citri and Poulsonii clades lack a CRISPR/Cas system, suggesting that it was absent in their common ancestor[39, 115]. However, our results from ccytper reveal a potentially functional CRISPR/Cas system in: (a) the Citri clade strain *s*Pte (<u>Supporting Figure S3</u>; <u>Supporting Dataset S6</u>), which is sister to the remaining members of Citri clade (Figure 2); (b) two Poulsonii clade strains (*s*Dum and *s*Ten) that are distantly related to those associated with *Drosophila* and the tsetse fly; and (c) two clade X strains. Although a functional CRISPR/Cas system is expected to prevent phage invasion/proliferation, all of these strains had at least two genes with a “phage” or “plectro” annotation. Similarly, as noted previously for TU-14 and NBRC_100390[39], several strains lacking CRISPR/Cas were mostly/completely devoid of “phage” and “plectro” gene annotations (Figure 2). The *Drosophila*-associated Citri and Poulsonii clade strains have very few “plectro” genes (range 0–3), whereas several non-*Drosophila*-associated strains had >20 (e.g. *s*Car, *S. phoeniceum*, *S. kunkelli*, *s*Rhe, *s*Ach, *S. citri*; Figure 2).

Further examination of sequences of potential plectroviral origin (i.e., Delta-blast search against the nr database using a “plectrovirus svts2 rep protein” as query), recovered genes found in *Spiroplasma* strains belonging to clades Ixodetis, clade X, Apis (only one strain), Poulsonii, and Citri. These sequences tended to show more similarity within clade (<u>Supporting Figure S4</u>; <u>Supporting Dataset S7</u>). No evidence of such proteins was found in the Citri clade strains *s*Ald-Tx, *s*Nigra, *s*Hy2, and *s*Moj (except for one short 54 aa ORF EHV01_0264). On the basis a the small number of genome assemblies at the time, Ku [116]hypothesized that susceptibility to plectroviral invasion originated in the common ancestor of the Citri+Poulsonii (as defined herein) or in one of its subclades. Our results are in line with that hypothesis, if several independent losses are assumed, but the presence of sequences of plectroviral origin in the Apis and Ixodetis clades (Supporting Figure S4) implies either additional independent invasion(s), or a single invasion in the ancestor of Citri+Poulsonii+Ixodetis+Apis, followed by multiple independent losses.

In addition to Plectroviridae[when active, characterized by non-enveloped rigid rods containing single-stranded DNA;117], other types of particles and/or DNA sequences/features derived from phage occur in *Spiroplasma[reviewed in* 118, 119]. Ramirez [118] assembled phage-like contigs (∼19kbp) from DNA isolated from phage-like particles in the Poulsonii clade strains *s*Mel and its close relative *s*Neb (a.k.a. “NSRO”; original host = *Drosophila nebulosa*). Regions with substantial homology to the *s*Mel phage-like contig are detected in most *s*Mel assemblies[118], suggesting that they might be lysogenic. We searched for the presence of such phage-like contigs in the *s*Ald-Tx, *s*Hy2 and *s*Moj genome assemblies by Geneious “Map to reference” and blastn of assembled contigs to the *s*Neb and *s*Mel phage-like sequences. The only result from this search was EAKCHEMG_367 (23,369 bp) in *s*Hy2, which is also the only contig obtained from the long reads (Nanopore) dataset. EAKCHEMG_367 contained several genes associated with phage genomes, such as portal, head-tail connector, capsid protein, recT, terminase (large and small subunits), and transposase-like (<u>Supporting Figure S5</u>), and had higher coverage (approximately 10 times higher than regions of contigs that are not repetitive, based on the short-read dataset; not shown). Nucleotide similarity between *s*Hy2’s EAKCHEMG_367 and the *s*Mel and *s*Neb phage-like contigs ranges ∼65–72% (<u>Supporting Figure S5</u>; alignment available in <u>Supporting Dataset S8</u>).

#### 3.4.2 Annotations based on Clusters of Orthologous Groups (COG) and the Kyoto Encyclopedia of Genes and Genomes (KEGG)

Based on the COG analysis of the three *Drosophila*-associated Citri clade strains, the most abundant category of genes was “Translation, ribosomal structure and biogenesis” (COG=J), followed by “Replication, recombination and repair” (COG= L) (<u>Supporting</u> <u>Figure S6</u>). This pattern was reversed (i.e., category L had the most genes, followed by J) in the Poulsonii clade strains (*s*Mel, *s*Hy1 and *s*Neo), but it appears to be driven by the large number of genes annotated as transposases, which fall under L (<u>Supporting</u> <u>Dataset S9</u>).

Respectively for *s*Ald-Tx, *s*Moj, and *s*Hy2, of the 914, 871, and 944 predicted protein-coding genes, 397, 362, and 397 could be assigned functional predictions in the form of KEGG Orthology (KO) numbers. A Venn diagram comparison of the genes assigned KO numbers indicates that the three strains share 336 genes (<u>Supporting Figure S7</u>; <u>Supporting Dataset S10</u>); these analyses only count one copy of each KO number per strain. *s*Ald-Tx and *s*Moj, which are each other’s closest sister (Figure 2) but whose hosts (*D. aldrichi*, *D. mulleri*, and *D. mojavensis*) are members of the *mulleri* complex, share the most KO genes (336 + 42; <u>Supporting Figure S7)</u>. The number of strain-unique KO genes ranged from 9 to 12. Based on Brite categories, the largest differences in KO gene content among the three strains are in the category of Enzymes (<u>Supporting Figure S8</u>; <u>Supporting Dataset S11</u>). For comparison, the three Poulsonii clade strains *s*Mel, *s*Hy1, and *s*Neo, respectively, had 635, 553, and 520 genes assigned KO numbers. Of these, 413 are shared and 4–9 are strain-unique <u>Supporting</u> <u>Figure S9</u>; <u>Supporting Dataset S12</u>).

Below we compare particular sets of genes among five strains of the Citri clade and five strains of the Poulsonii clade. For the Citri clade, we included the *Drosophila*-associated *s*Ald-Tx, *s*Hy2, *s*Moj, as well as *s*Nigra (host: non-*Drosophila* Dipteran), and *S. melliferum* (a culturable and horizontally transmitted symbiont of the honey bee that has comparatively greater metabolic capacities). For the Poulsonii clade, we include the reference genome for each of the three available *Drosophila*-associated strains (*s*Mel, *s*Hy1, and *s*Neo), their sister lineage *s*Car (from the flesh fly *Sarcophaga carnaria*), and a more distant relative (*s*Fus) associated with the tsetse-fly (*Glossina fuscipes fuscipes*).

#### 3.4.3 DNA Repair Related Genes

Within the Homologous Recombination and the Base Excision Repair pathways, the *Drosophila*-associated Citri clade strains tend to have more missing genes than their Poulsonii clade counterparts (<u>Supporting Table S7</u>). Previous research on the Citri clade strains *S. melliferum* and *S. citri* indicate they are RecA-deficient, rendering them highly sensitive to UV radiation[120]. *RecA* was frame-shifted, incomplete or absent in the five Citri clade genomes compared (<u>Supporting Table S7</u>), and appears to be lacking in all the Citri clade assemblies (not shown). In contrast, all Poulsonii clade except *s*Mel [121] and sCru, appear to have a functional copy of *RecA* (not shown). Little to no difference in gene content is detected among the ten strains compared regarding the Nucleotide Excision Repair and the Mismatch Repair pathways. A notable difference between the two clades is that the five Citri clade strains compared in <u>Supporting Table S7</u> encode Deoxyribodipyrimidine photo-lyase (*phrB*), whereas all five Poulsonii clade strains do not. Nonetheless, *phrB* is encoded by Poulsonii clade strains TU-14 and S. sp. NBRC_100390 (not shown), the closest known relatives of *s*Fus (Figure 2), suggesting independent losses in *s*Fus and in the ancestor of *Poulsoni* s.s. In general terms, it appears that the *Drosophila*-associated Citri clade strains have similar or worse abilities to repair DNA, and are thus likely to evolve equally or more rapidly, than their Poulsonii clade counterparts, which have among the highest DNA substitution rates of bacteria[39].

#### 3.4.4 Inferred abilities to import and process certain metabolites

The ability to import and process molecules associated with energy metabolism appears to be substantially limited in the *Drosophila*-associated Citri clade strains (*s*Ald-Tx, *s*Hy2, *s*Moj; summarized in <u>Supporting Figure S10</u>). As detailed below, the annotation suggests that the only sugar they are able to import (and metabolize) is fructose. Comparatively, at least one of the three members of the *Drosophila*-associated Poulsonii clade are predicted to be able to convert pyruvate to acetyl-CoA (pyruvate oxidation; Figure 2), and to import and process glucose[functionally confirmed in sMel; 121], fructose (except *s*Mel), glycerol (which is predicted to produce peroxide), and arginine (only *s*Hy1).

Of the ten strains compared in <u>Supporting Table S8</u>, the Citri clade *S. melliferum* and *s*Nigra, and the Poulsonii clade *s*Hy1, *s*Mel, *s*Neo, and *s*Fus are predicted to be able to import and process glucose (i.e., they have putatively functional *ptsG*, *crr*, and *pgi* genes). In contrast, the remnants of at least two of these three genes appear non-functional in the *Drosophila*-associated Citri clade strains (*s*Ald-Tx, *s*Hy2 and *s*Moj) and in the Poulsonii clade *s*Car, suggesting they cannot use glucose as an energy source, unless they can import glucose employing other putative sugar transporters encoded by their genomes (e.g. locus tags EHU54_02420, EHU54_02680, EHV01_02065, EHV01_02360, EHV01_01190, BST80_02715, BST80_01055). All Citri and Poulsonii strains compared, except for *s*Mel, are predicted to be able to import and metabolize fructose, with all strains except *s*Mel, *s*Hy1 and *s*Neo encoding two putatively functional *fruA* genes (<u>Supporting Figure S10</u>; <u>Supporting Table S8</u>); *s*Neo and *s*Hy1 retain one functional *fruA* and non-functional remnants of another putative *fruA* gene. Of the ten strains compared, only *s*Hy2 (EHV01_02360) and *S. melliferum* have a putatively functional importer of trehalose (*treB*), but all strains compared except *S. melliferum* lack a functional *treA* gene (the few other *Spiroplasma* strains that appear to encode *treA* and/or *treB* are listed in <u>Supporting Dataset S13)</u>. Therefore, none of the Diptera-associated strains we compared in <u>Supporting Table S8</u> seem to be able to use trehalose, the main sugar in insect hemolymph[reviewed in 122]. Of the ten strains compared, only *S. melliferum* encodes all four genes needed for uptake and metabolism of N-Acetylmuramic acid (i.e., *murP*, *murQ*, *nagA*, n*agB*). The ability to import and metabolize glycerol or produce H_2_O_2_ (based on the presence of genes encoding *glpF*, *glpK*, and *glpO*) appears to be restricted to *S. melliferum* (Citri clade), and all but one (*s*Fus) of the five Poulsonii clade strains compared.

Of the ten strains compared, only *s*Hy1 and *S. melliferum* appear to encode the machinery needed to use import and metabolize arginine (i.e., the adjacent genes *ArcA*, *ArcF*, *ArcC*, *ArcD*) (<u>Supporting Figure S10</u>; <u>Supporting Table S8</u>). Of the ten strains compared, only the *Drosophila*-associated Citri Clade (*s*Ald-Tx, *s*Hy2, and *s*Moj) lack the full operon associated with with pyruvate oxidation (pyruvate => acetyl-CoA; Koala M00307; genes *pdhA*, *pdhB*, *pdhC*, *pdhD*) (<u>Supporting Table S8</u>; Figure 2). All the strains compared except for *S. melliferum* lack functional copies of the five genes needed to import/process cellobiose (<u>Supporting Figure S10</u>; <u>Supporting Table S8</u>). *GlpX* (K02446, which catalyzes GA-P3 >> Fructose-6-P) appears to be missing from the five Poulsonii strains compared (<u>Supporting Table S8</u>), but it is present in NBRC_100390 and TU-14 (WP_070407118.1). Of the five Citri clade strains compared, *GlpX* is present in *S. melliferum*, *s*Nigra and *s*Moj, but it is pseudogenized in *s*Ald-Tx and *s*Hy2 (<u>Supporting Table S8</u>; Figure 2).

The *Drosophila*-associated Citri clade strains (*s*Ald-Tx, *s*Hy2, *s*Moj) appear to be more limited than their Poulsonii-clade counterparts (*s*Mel and *s*Hy1) regarding the ability to generate phospholipids from fatty acids or diacylglycerol (DAG). Each of the *Drosophila*-associated Citri clade strains lack at least one functional gene of the terpenoid C55 synthesis pathway (<u>Supporting Table S8</u>). Similarly, at least one of the four genes needed to convert DAG-3P to cardiolipin is missing or pseudogenized in *s*Ald-Tx, *s*Hy2, *s*Moj, whereas *s*Mel [121] and *s*Hy1 are predicted to have this capacity (<u>Supporting</u> <u>Table S8</u>}.

The three *Drosophila*-associated Citri clade strains lack a functional copy of ferritin-like genes (*Ftn*), which are present in the remaining seven strains compared in <u>Supporting</u> <u>Table S8</u>. Ferritin-like proteins are involved in iron sequestration[123]. A search for the term “ferritin” in the gene annotations of the recently released genome assemblies of the Citri and Poulsonii Clade indicate three additional strains lacking this gene, which appear to have independently lost it (i.e., *s*Pub, *s*Cru, and *s*Ten; not shown). Two observations suggest that an *Ftn* gene is important for the *sMel*-*D. melanogaster* symbiosis: higher *Ftn* transcript levels inside the host compared to in vitro culture[23]; and *Ftn* is one of few genes whose protein abundance is upwardly biased compared to transcript levels[42]. Iron homeostasis is relevant to insect-symbiont associations[e.g. 124], including that of *Drosophila* and *Spiroplasma.* Strain *s*Mel induces expression of *Drosophila* transferrin gene *Tsf1*, which binds and facilitates the sequestration of iron from the hemolymph to the fat body[35, 125]. Proliferation of *Spiroplasma* in *D. melanogaster* (both *s*Mel and the plant pathogen *S. citri*) requires Tsf1-bound iron[125]. The lower levels of free iron in the hemolymph appear to underlie the *s*Mel-induced resistance against two pathogens[the bacterium Providencia alcalifaciens and the fungus Rhyzopus oryzae; 35]. If the *Spiroplasma*-encoded *Ftn* genes are involved in the ability of *Spiroplasma* to exploit iron, the *Drosophila*-associated Citri clade strains (*s*Ald-Tx, *s*Hy2, *s*Moj) likely have limited iron exploitation capacities.

Therefore, overall the *Drosophila*-associated Citri clade strains (*s*Ald-Tx, *s*Hy2, *s*Moj) seem metabolically more limited than their Poulsonii clade counterparts (*s*Mel, *s*Hy1, and *s*Neo) regarding the importation and processing of several metabolites including sugars, lipids, and iron. The inability to exploit such resources may underlie their comparatively lower densities and vertical transmission rates in native and non-native hosts[38, 98, 126; and personal observation].

#### 3.4.5 Putative virulence factors

The genomes of *Spiroplasma* encode a diverse set of known or putative toxins genes, some of which have been mechanistically linked to phenotypes, such as male killing and parasite killing[19, 43, 46, 47, 51, 73, 93, 107, 109, 121, 127–130]. Among these, genes with domains annotated as Ribosome Inactivating Proteins (RIPs) appear to be the most common and diverse, and are predominantly found in vertically transmitted *Spiroplasma*, based on a comparison of 12 vertically- and 31 horizontally-transmitted *Spiroplasma* genomes available at that time[19]. RIPs include toxins such as ricin (from the castor oil plant) and Shiga toxin (from *E. coli*). RIPs target an adenine found within a 12 nucleotide motif of the 28S rRNA that is universally conserved in eukaryotes, termed the sarcin-ricin loop. RIPs remove the target adenine, leaving an abasic (a.k.a. depurinated) site, which irreversibly renders the ribosome non-functional and thus stalls protein synthesis[reviewed in 131]. The toxicity of RIPs varies based on their ability to enter the cell, to reach the appropriate cellular compartment, and to resist degradation[132]. Evidence that the presence of *Spiroplasma* induces ribosome depurination exists for two *Drosophila*-associated Poulsonii clade strains. Presence of *s*Neo in its *Drosophila* host confers protection against nematodes and wasps; both macroparasites exhibit signals of ribosome depurination in the presence of *s*Neo; and a recombinantly produced RIP protein encoded by sNeo (i.e., *s*Neo_RIP1_WP_127093322; <u>Supporting Figure S11</u>; <u>Supporting Dataset S14</u>) has confirmed RIP activity, as it depurinates the target adenine in vitro of both whole nematode and cell-free rabbit ribosomes[43, 93]. Similarly, presence of *s*Mel in its *Drosophila* host confers protection against certain wasps; such susceptible wasps exhibit signs of ribosome depurination, depurination of *Drosophila* ribosomes is also detected (at least at the embryo stage); and ectopic expression of two *s*Mel encoded RIP genes (i.e., MSRO_RIP1_WP_040093770 and MSRO_RIP2_WP_040093936; <u>Supporting Figure S11</u>) confirms their RIP activity against host ribosomes[51, 87, 88, 92, 93, 128, 129].

Our results reveal the presence of domains identified as RIPs in numerous additional *Spiroplasma* strains, including the three *Drosophila*-associated Citri clade strains: *s*Moj (1 gene), *s*Ald-Tx (2 genes), and *s*Hy2 (4 genes) (Figure 2; <u>Supporting Figure S11</u>). All but five Citri clade strains encode at least one RIP gene, including several first records (*s*Pte, *s*Nob, *s*Chi, *s*Pub, and *s*Tum). All but one Poulsonii Clade strains (i.e., *s*Dum) have at least one RIP gene, including several first records (*s*Car, TU-14, NBRC_100390, *s*Fus, *s*Cru, *s*Ten, *s*Cal). Similarly, in the newly identified Clade X, all but one strain had at least one RIP gene. Several new RIP gene records were found in the Ixodetis and Apis clades (<u>Supporting Figure S11</u>).

Although we recovered 175 RIP domain sequences, 31 sequences were excluded from phylogenetic analyses because they were very short (<u>Supporting Dataset S14</u>). Many nodes in the RIP domain amino acid phylogeny received low support by one or more of the clade support measures (<u>Supporting Figure S11</u>). While there are a few RIP domain clades that are restricted to the same *Spiroplasma* strain or *Spiroplasma* clade, there are several cases where RIP domains recovered as sister lineages belong to distantly related *Spiroplasma* strains. For example, the single *s*Moj RIP gene (*s*Moj_RIP1_MBH8624287) appears closely related to RIPs from Clade X (*s*Lun), Citri (*s*Had), and Poulsonii (*s*Hy1 WP_198049692 and *s*Mel WP_040093770). One RIP from *s*Hy2 (*s*Hy2_EHV01_04225) was recovered as sister to a RIP from Clade X (*s*Cur_WP_342223867), and was closely related to RIPs from other Clade X (sLute_WP_338968587 and sLun_WP_338981923) and from Poulsonii (sNeo_RIP4_WP_158676203). One RIP from *s*Ald-Tx (*s*Ald_Tx_EHU54_01310) appears as sister to the RIP of *s*Cerv (Citri), with their next most closely related RIPs belonging to Clade X (*s*Flav, *s*Cur, *s*Lute).

The remaining RIP-encoding genes from *s*Hy2 (3 genes) and *s*Ald-Tx (1 gene) fell within a well supported clade (“Clade RAO”; <u>Supporting Figure S11</u>). Most of the genes in this clade have an unusual domain structure, previously referred to as “Spaid-like” by Moore and Ballinger [19] and “RIP/Spaid” byFilée [109], containing two RIP domains, followed by Ankyrin repeats (A); an OTU domain (ovarian tumor deubiquitinase; IPR003323; O), and a C-terminal hydrophobic (transmembrane) domain (Figure 3). Clade RAO is composed predominantly of genes from clades X (first record) and Citri, as well one strain from each of three additional *Spiroplasma* clades: Ixodetis (*s*Chrys WP_174481319; not shown); “sister to Ixodetis” (the *s*Riversi WP_215825920 <u>Supporting Figure S11</u> and WP_215826391; not shown); and Poulsonii. The Poulsonii clade RAO gene (*s*Hy1_RIP2_MBH8623170) appears most closely related to *s*Hy2_EHV01_02330 (Citri), suggesting a horizontal transfer between these distantly related strains that overlap geographically and share the same host species[27, 30, 31, 52]. Another *s*Hy2 RIP (*s*Hy2_EHV01_01950) is recovered as closely related to RIPs from several Citri Clade strains associated with Lepidoptera, and Clade X strains associated with Diptera and Coleoptera.

**Figure 3.**
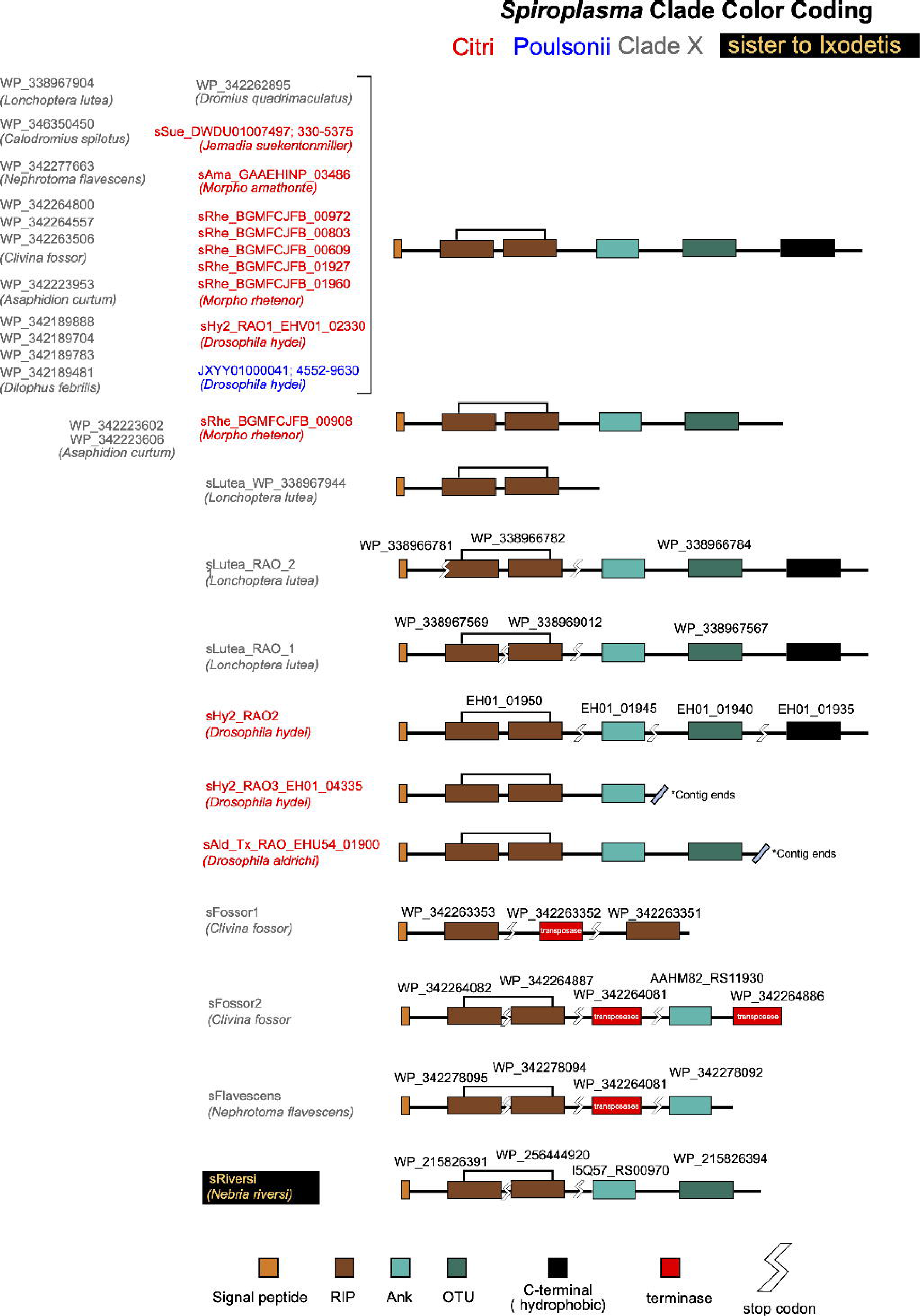
Domain architecture (not to scale) of genes in the clade RAO (RIP, Ankyrin, OTU).

To further explore potential patterns of HGT, we examined the DNA sequences of clade RAO genes, including adjacent regions in cases where a stop codon or transposase gene interrupts domains. We arbitrarily broke up the alignment into 5 blocks that were separated by several poorly conserved regions (based on visual examination of the identity graph in Geneious; <u>Supporting Dataset S15</u>). We performed phylogenetic analyses on each of these 5 regions, which revealed patterns of topological incongruence among regions (<u>Supporting Figure S12</u>). We highlight for example that in the first block (N-terminus), *s*Hy2_EHV01_02330 appears as most closely related to RAO in *s*Hy1, consistent with the amino acid phylogeny of only the RIP domain (<u>Supporting Figure S11</u>). In contrast, the Ankyrin Repeats Region of *s*Hy2_EHV01_02330 appears most closely related to two Clade X RAOs from strain sFossor (WP_342264303 and WP_34226455). These observations are consistent with the features of polymorphic toxins; i.e., multidomain secreted proteins that diversify through recombination and domain swapping, and tend to be associated with horizontally transferred elements[73, 133].

OTU domains, which are common in eukaryotic genes, with and without RIP and/or Ankyrin domains have previously been reported in genes of several *Spiroplasma* strains[e.g. 19, 73, 107, 109, 121, 127], including in *Spaid[*Ankyrin + OTU male-killing gene of sMel; 47]. According to the analysis ofMoore and Ballinger [19], within *Spiroplasma*, OTU and Ankyrin repeat domains are exclusive to vertically transmitted strains. In addition to OTU domains present in “RAO” genes, we detected OTU domains in several additional Citri clade strains (e.g. *s*Moj, *s*Tum, *s*Man) and Clade X (Figure 2). Within the Poulsonii clade, however, OTU domains are only detected in the three *Drosophila*-associated strains (Figure 2), which are known to be vertically transmitted. Within the Citri clade, it is notable that OTU domains are absent in the plant-associates (e.g. *S. phoeniceum*, *S. kunkelii*, *S. citri*, and *S. sp.* Chis) and the horizontally-transmitted pathogen of honeybees (*S. melliferum*). All other Citri clade strains have at least one OTU domain-containing gene and are associated with insects (Diptera, Lepidoptera, Hymenoptera, and one Coleoptera; Figure 2). The transmission mode of most of these newly available strains is unknown (most were discovered in the process of assembling and annotating their host’s genome sequence), but given the presence of OTU and Ankyrin repeat domains, they are likely vertically transmitted. The OTU domain tree recovers a clade comprised exclusively of all but four of the OTU-containing genes from *Drosophila*-associated strains (both Poulsonii and Citri clades; Supporting Figure 13; Supporting Dataset S16), including *Spaid* (WP_105629072), which could indicate a common origin or function. The exceptions are: *s*Hy1dLiv_PMBNAAIA_00948_AO (sister to an OTU from a Citri clade *Myrmica* ant associate); *s*Hy2_EHV01_02330_RAO (sister to Citri clade strains associated with Lepidoptera); *s*Hy2_EHV01_01940 (sister to a Citri clade strain associated with a *Morpho* butterfly); and sAld_EHU54_01900_RAO (sister, albeit with low support, to two genes from a Clade X strain, and embedded within a well-supported clade that includes Citri clade strains from Lepidoptera and Coleoptera). In contrast, the RIP domain of *s*Ald_EHU54_01900_RAO is closely related to genes from *s*Hy1 (Poulsonii), *s*Hy2 and *s*Teh (Citri clade) (Supporting Figure S11), suggesting that *s*Ald_EHU54_01900 is the result of horizontal transfer and domain shuffling.

Strains *s*Ald-Tx and *s*Hy2 encode a gene containing an OTU domain and another toxin domain (i.e., ETX/MTX2; β-pore-forming toxins with receptor-binding activity; hereafter “ETX”); an unusual domain architecture that was not found in any other *Spiroplasma* except *s*Hy1[sHy1_00057; 39]. Genes encoding components of the ETX pore-forming protein have been reported in several *Spiroplasma* genomes [reviewed in 73] and tend to be associated with vertically transmitted strains[19]. We detected the ETX domain in additional strains from the following clades: several from Poulsonii; all but two strains of Citri (only absent in the plant-associates *S. phoeniceum* and *S. kunkelii)*; Mirum; all but one of Clade X strains (Figure 2); Apis; and Ixodetis (Supporting Figure S14). The ETX domain of *s*Ald-Tx, *s*Hy2 and the Poulsonii-clade *s*Hy1 were highly similar, including adjacent regions Supporting Figure S15 (Supporting Dataset S18) and grouped together (Supporting Figure S14; Supporting Dataset S17), implying a recent horizontal transfer between the Poulsonii and Citri clade. These were in turn sister to a clade composed of genes in other strains of the Citri clade, as well as strains in Clade X, and one strain “sister to ixodetis”, collectively associated with Diptera, Coleoptera, and Hymenoptera. The ETX domain-containing gene of *s*Moj (BST80_00145 = MBH8623564.1) was most similar to that of a Clade X strain associated with a beetle (WP_342264206.1). With the exception of ETX genes in the Mirum clade, there is a general lack of monophyly of ETX domains within *Spiroplasma* strains and clades, suggestive of substantial losses and gains from distantly related strains.

A gene encoding an ETX domain along with an upstream PQQ-like domain (PFAM: PF13570), and an N-terminal Signal Peptide is found in the Poulsonii clade strain *s*Neo (WP_127092276) and the Citri clade strain *s*Nigra (WP_126821430), both from Dipteran hosts associated with mushrooms. We found evidence of this PQQ-ETX domain architecture in one Clade X strain associated with a beetle, and in six additional Citri clade strains (*s*Chi, *s*Rhe, *s*Ach, *s*Ama, *s*Tum, and *s*Man), but not in those associated with *Drosophila* (Figure 2). Additional strains encoding a PQQ domain-containing gene, but lacking an ETX domain, were detected in clades Poulsonii, Citri, Chrysopicola, Mirum, and X (Figure 2). In *s*Mel (Poulsonii), the neighboring CDS WP_040094248.1 (PQQ) and WP_040094250.1 (ETX) appear to reflect the remnant of a PQQ-ETX protein. A phylogeny of the PQQ domain alignment (<u>Supporting Figure</u> <u>S16</u>; <u>Supporting Dataset S19</u>) had very poor resolution and is thus not shown. The function of PQQ-domain-containing proteins in *Spiroplasma* is unknown. PQQ-like enzymes have repeats of a ß propeller. Depending on the number of blades they contain, ß propellers have a diversity of functions, including ligand binding, transferase, hydrolase, lyase, isomerase, signaling, oxidoreductase, and structural protein[134].

The widespread occurrence in *Spiroplasma* of genes with the RAO domain architecture, whose RIP domain region forms a monophyletic group, implying a common origin, is intriguing. Ankyrin repeats are a common structural motif (33 aa) involved in protein-protein interactions. They are ubiquitous in eukaryotic proteins, but also found in viruses, archaea, and they are particularly common in bacteria with a symbiotic (pathogenic to mutualistic) lifestyle[135]. In *Spaid*, the male-killing gene of strain *s*Mel, the ankyrin repeats domain is required for Spaid to accumulate on the male X chromosome, a prerequisite for its male killing action[47]. Similarly, in the human pathogen *Legionella pneumophila*, the ankyrin repeats domain of a particular protein is used to specifically bind to a host protein on which damage (i.e., phosphocholination) is exerted by another domain of the same protein[136]. Therefore, we hypothesize that through its protein-binding function, the ankyrin repeats domain of RAO and other *Spiroplasma* genes enables localization to specific eukaryotic cells or subcellular regions. Because of the apparent role of at least some *Spiroplasma* RIPs in host defense, ankyrin repeats might enable the selective localization of RIP (whose ribosomal RNA target motif is universally conserved in eukaryotes) and its depurination activity, to the cell or cell compartments of the endo-macroparasite (e.g. parasitic wasp or nematode), rather than to those of the host. Regarding the potential role of OTU in RAO, the recent work on *Spaid* by Harumoto [46] offers potential clues. OTUs encode deubiquitinase activity, which reverses protein ubiquitination, an important post-translational modification in eukaryotes, that influences stability, interactions, activity and localization of proteins. Harumoto [46]demonstrated that: (1) *Spaid* lacking the OTU domain exhibits attenuation of the male killing action, because the OTU-free Spaid protein is polyubiquitinated and degraded via the host ubiquitin-proteasome pathway; and (2) Spaid uses its OTU domain to deubiquitinate itself in an intermolecular fashion, presumably through a homomeric interaction. Consequently, the OTU domain functions as a self-stabilizing mechanism for Spaid. We hypothesize that the OTU domain in RAO and other *Spiroplasma* genes could serve such a self-stabilizing role.

Our results indicate that the genomes of the *Drosophila*-associated Citri clade strains *s*Moj, *s*Ald-Tx, and *s*Hy2 encode a putative toxin that appears to be exclusive to the Citri clade: ADP-ribosyltransferase exoenzyme; PFAM:PF03496; or lethal factor; <u>Supporting</u> <u>Figure S17</u>; <u>Supporting Dataset S20</u>). ADPs convert nicotinamide adenine dinucleotide (NAD) into nicotinamide and ADPribose (ADPr), which is transferred to proteins, nucleic acids, or small molecules. Such modifications can inhibit normal function of host proteins or activate host proteins in a manner that promotes bacterial fitness[137–140]. ADP is typically part of a binary toxin that also includes the protective antigen (PA) protein that enables entry into the host’s cell[reviewed in 137]. Of all *Spiroplasma* sequenced to date, only *S. melliferum* encodes putative functional genes for both components (<u>Supporting Figure S17</u>), but it was not found to cause the expected cytotoxic effect in the yeast growth-deficiency assay, suggesting it requires a host-specific factor for activation[138]. Ten additional Citri clade species/strains encode remnants of one or both genes (<u>Supporting Figure S17</u>), all of which occur immediately downstream of two apparently functional CDS (not shown): an NAD(+)/NADH kinase (GO function: 0003951; GO_process: GO:0006741) and a PTS transporter subunit EIIB (GO function: 0008982). The three *Drosophila*-associated Citri clade strains encode a full-length ADP (annotated as “lethal factor CDS” in <u>Supporting Figure S17</u>). However, the PA gene is interrupted by an early stop codon (only retaining one of the three domains; the Ca-binding domain pfam 03495). Immediately downstream of the stop codon, *s*Moj has a remnant of a transposase (BST80_01070; PF13613), whereas in *s*Ald-Tx and *s*Hy2 the end of the contig is reached, likely reflecting the presence of a repetitive sequence that hampered assembly of this region. Remnants of PA’s domain 2 (pfam17475) and domain 3 (pfam 17476) are found in some lineages (<u>Supporting</u> <u>Figure S17</u>). The *Spiroplasma* ADP proteins resemble the C3-like ADP-ribosyltransferases, as evidenced by the conservation of two residue sites essential for enzymatic activity: the R site and the QXE motif (<u>Supporting Figure S17</u>). No evidence of signal or transport motifs exists in any of the *Spiroplasma* ADP-domain containing genes, raising doubts about their potential function, unless they function within the *Spiroplasma* cell. All of the *Spiroplasma* ADP genes share high homology with each other, and are very distinct from their closest blast hits (i.e., from *Bacillus*; not shown).

Among *Spiroplasma* proteins considered important for the interaction with the (insect) host are the abundant membrane lectins referred to as spiralins[141–143]. Each of the three *Drosophila*-associated Citri clade strains (*s*Ald-Tx, *s*Moj, and *s*Hy2) encode one gene annotated as the lipoprotein spiralin (hereafter *spiA*; <u>Supporting Table S8</u>). These spiralin gene sequences formed a monophyletic group, whose closest relatives are spiralin genes from other members of the Citri clade (<u>Supporting Figure S18 Spiralin</u>; <u>Supporting Dataset S21</u>), including a gene known to be required for efficient transmission by the insect vector to the host plant of *S. citri*[142]. In contrast, the six Poulsonii clade strains, including *s*Fus from *Glossina*, encode at least two highly divergent spiralin genes (<u>Supporting Figure S18</u>), identified in *s*Mel as *spiB* [121] and *spiC*[23]. In addition, all Poulsonii clade strains except *s*Fus, encode a spiralin gene that is more similar to those of the Citri clade (i.e., *spiA*). The *spiB* gene is the most highly expressed gene in *s*Mel[23, 121], and it appears to be involved in the process of vertical transmission during which *s*Mel enters the oocyte from the hemolymph[143]. It is possible that lacking a *spiB* or *spiC* homologue, contributes to the lower vertical transmission efficiency of the *Drosophila*-associated Citri clade strains.

## 4 Conclusions

Based on the draft assemblies contributed here, we infer that compared to their Poulsonii clade counterparts (*s*Mel, *s*Hy1, and *s*Neo), the *Drosophila*-associated Citri clade strains (*s*Ald-Tx, *s*Hy2, and *s*Moj) have smaller genomes, fewer genes, more limited metabolic capacities (including importation and metabolism of sugars, lipids, and iron), possibly lower ability to transmit vertically due to the absence of the *spiB* gene, and equally bad or worse DNA repair mechanisms. Collectively the above features may underlie their comparatively lower densities and vertical transmission rates, and lead us to predict that the *Drosophila*-associated Citri clade evolves at rates similarly high to those reported in the Poulsonii clade. Frequent loss and rapid evolution of symbionts, along with the lack of *in vitro* culture protocols and genetic tractability, pose practical challenges to experiments assessing phenotypes and their underlying genetic basis. Notwithstanding their comparatively “poor” features, the genomes of *s*Ald-Tx, *s*Hy2, and *s*Moj collectively encode a similarly diverse repertoire of putative toxin genes to the Poulsonii clade, with evidence of substantial exchanges, including between the Poulsonii and Citri clade (e.g. the RAO gene of *s*Hy1 and *s*Hy2), and within-genome shuffling. Presumably some of these toxins are used in the interaction with their *Drosophila* host or their host’s natural enemies, but based on examination of two strains (*s*Ald-Tx and *s*Moj) in the presence/absence of two different parasitic wasps (including one highly susceptible to Poulsonii clade strains), no evidence of fitness consequences to their native hosts has been detected. How Citri clade strains persist, and even achieve high prevalence, in (certain) *Drosophila* populations remains a mystery. It is possible that they confer yet undiscovered net fitness benefits to their (female) hosts or that they rely on substantial horizontal transmission. Beyond the *Drosophila*-associated Citri clade, the discovery of a divergent *Spiroplasma* lineage associated with dipterans and coleopterans (clade X) underscores the cryptic diversity of endosymbionts that metazoan genome projects are uncovering. The common occurrence of RAO domain architecture in very distant *Spiroplasma* lineages (predominantly in clades X and Citri) suggests that it plays an important role in insect-*Spiroplasma* interactions, which may include a combination ribosomal RNA depurination (by RIP), selective localization to target cells or cell compartments (by Ankyrin repeat proteins), and self-stabilization (by the OTU deubiquitinase).

## Supporting information

Supporting Protocol S3

Supporting Protocol S1

Supporting Protocol S2

## Author Contributions

Designed and conceived project: PR, HMM, MM, RA

Fitness experiments: HMM, MM

Insect husbandry: PR, HMM, MM

Symbiont DNA purification and sequencing: PR, HMM

Assembly and annotation: PR, HMM, MM, RA

Phylogenetic analyses: PR, MM, HMM

Wrote first draft of paper: MM, PR, HMM

All authors contributed to revising the manuscript, and have read and approved the final manuscript.

## Acknowledgements

Caitlyn Winter and Lauryn Winter collected the *D. mojavensis* line at Catalina Island. Joyce Kao and Sergey Nuzhdin organized and led the Catalina Island field trip. The Wrigley Marine Science Center provided housing, island transportation, and access to their research facilities on Catalina Island. Texas State University at San Marcos granted access to Freeman Ranch. The University of Texas at Austin granted access to Brackenridge Field Laboratory. Esperanza Martinez-Romero (Centro de Ciencias Genómicas, UNAM, Mexico) hosted MM during establishment of *D. hydei* line. Jordan Jones provided statistical advice. Igor Vilchez Ramirez contributed preliminary data regarding *repleta* group flies in the Texas Hill Country. Todd Schlenke provided *Leptopilina heteroma* strain Lh14.

Portions of this research were conducted with the advanced computing resources provided by Texas A&M High Performance Research Computing. The Galaxy server that was used for some calculations is in part funded by Collaborative Research Centre 992 Medical Epigenetics (DFG grant SFB 992/1 2012) and German Federal Ministry of Education and Research (BMBF grants 031 A538A/A538C RBC, 031L0101B/031L0101C de.NBI-epi, 031L0106 de.STAIR (de.NBI)). For the use of the *Glossina fuscipes fuscipes* sequence assembly, we acknowledge Dr. Serap Aksoy and The Genome Institute, Washington University School of Medicine.

## Supporting Command Lines

**Supporting Command Line S1.** Command Line used to perform statistical analyses and graphs in R for the *Drosophila mojavensis Spiroplasma* protection experiments. 10.6084/m9.figshare.c.7437997

## Supporting Tables Descriptions

**Supporting Table S1.** Insect species and strains/isofemale lines used for genome assembly and/or fitness assays.

**Supporting Table S2.** Restriction enzymes and conditions to differentiate Spiroplasma strains sHy1 (poulsonii clade) and sHy2 (citri clade).

**Supporting Table S3.** List of Spiroplasma strains used for a local Blastn search. 10.6084/m9.figshare.c.7437997

**Supporting Table S4.** List of *Glossina fuscipes fuscipes* contigs assigned to *Spiroplasma*. 10.6084/m9.figshare.c.7437997

**Supporting Table S5.** *Spiroplasma* frequencies in wild-caught *D. aldrichi* and *D. mulleri* from sampled localities of Texas.

**Supporting Table S6.** P-values and assumed distribution (binomial or quasibinomial) from Analysis of Deviance Table (Type II tests) for the experiments that tested whether *Spiroplasma s*Moj influenced the interaction between *Drosophila mojavensis* and two species of parasitic wasps. None of the p-values were significant after Bonferroni correction for multiple comparisons (adjusted alfa = 0.05/11 = 0.0045).

**Supporting Table S7.** DNA Repair associated genes that are present in at least one of the insect endosymbionts *Arsenophonus nasonae*, *Wolbachia* (*w*Mel), and *Buchnera aphidicola* (APS). Comparison of annotations (from BlastKoala and/or NCBI) for ten *Spiroplasma* strains (red font = Citri clade; blue font = Poulsonii clade) for genes associated with DNA repair. Colored cells (yellow or red imply nonfunctional or absent gene). Locus tags or Protein IDs (WP) are provided for a subset of strains (mostly the newest ones).

**Supporting Table S8.** Comparison of genes associated with energy metabolism, ferritin and spiralin among ten strains of the Citri (red headers) and Poulsonii (blue headers) clades. Green cells = we infer that the gene is functional. Yellow/red cells = gene is absent or inferred as non-functional (e.g. too short). Locus tags and or protein IDs are given for some. Files used to identify genomes encoding (pseudo)gene for “1-acyl-sn-glycerol-3-phosphate acyltransferase” and Alpha/beta hydrolase or lysophosphatase, lysophosphatase” of “alpha/beta hydrolase” are found in <u>Supporting Dataset S22</u>.

## Supporting Figure Legends (all Supporting Figures available at 10.6084/m9.figshare.c.7437997)

**Supporting Figure S1.** Neighbor-Joining Tree based on Jukes Cantor (JC) distances among the 16S rDNA genes of many *Spiroplasma* strains, rooted with *Mycoplasma pneumoniae*. We provide a preliminary assignment of most strains to major clades (labeled and color coded). “Clade X” had not been reported before. Detailed phylogenomic analyses of the Citri and Poulsonii clades (with Mirum and Chrysopicola) are shown in Figure 2.

**Supporting Figure S2.** Strict consensus of 5 most parsimonious trees of the 16S rDNA gene of representative *Spiroplasma* strains in the Citri clade and one representative of the Poulsonii clade for rooting purposes.

**Supporting Figure S3.** Plot of CRISPR-Cas loci, orphan Cas operons, and orphan CRISPR arrays generated by https://crisprcastyper.crispr.dk/. Interference module in yellow. Adaptation module in blue. Accessory genes in purple. Cas6 in red. Arrays with their associated subtype in black/white checkerboard. Unknown genes in grey. Cas genes with low-quality alignments in lighter color with parentheses around the name. For circular topology, black vertical line denotes start/end of sequence. Numbers around arrays are (false) ORFs overlapping with predicted CRISPR arrays.

**Supporting Figure S4.** Neighbor-Joining (NJ) tree based on Jukes-Cantor (JC) distance among the subset of longest representatives of sequences assigned to plectrovirus.

**Supporting Figure S5.** A. Alignment (MAFFT default parameters) of the sHy2 (Citri clade) putative phage contig with those of *s*Neb and *s*Mel (Poulsonii clade) described in Ramirez [118]. Black and grey, respectively, represent sequence different/identical to consensus (0% majority). White block arrow = repeated sequences in *s*Hy2 (percent identity between repeats is indicated). Colored block arrows are inferred coding sequences (CDS). Yellow = hypothetical CDS with no homology to known proteins with known functions. Blue = transposase proteins. Green = capsid structure proteins. Orange = DNA-binding/recombination proteins. HJS = Holliday junction resolvase. sml. = terminase small subunit. For further details see Ramirez [118]. B. Percent nucleotide identity based on above alignment.

**Supporting Figure S6.** Number of genes by Cluster of Orthologous Groups (COG) category in the three *Drosophila*-associated *Spiroplasma* strains *s*Ald-Tx, *s*Hy2, and *s*Moj.

**Supporting Figure S7**. Venn diagram comparing shared and unique genes, according to KO numbers, among the *Drosophila*-associated Citri clade strains *s*Ald-Tx, *s*Moj, and *s*Hy2. Only one gene assigned per KO number per strain is used for these analyses. Venn diagrams generated with https://bioinformatics.psb.ugent.be/webtools/Venn/ and edited in Inkscape.

**Supporting Figure S8** Number of KEGG Orthology genes present and absent in the pairwise comparisons of *s*Ald-Tx, *s*Moj, and *s*Hy2. Brite descriptions (and families in grey ovals) are indicated.

**Supporting Figure S9.** Venn diagram comparing shared and unique genes, according to KO numbers, among the *Drosophila*-associated Poulsonii clade strains *s*Mel, *s*Hy1, and *s*Neb. Only one gene assigned per KO number per strain is used for these analyses. Venn diagrams generated with https://bioinformatics.psb.ugent.be/webtools/Venn/ and edited in Inkscape.

**Supporting Figure S10.** Predicted sugar metabolism capabilities of the three *Drosophila*-associated *Spiroplasma* strains belonging to the Citri clade (*s*Moj, *s*Hy2, and *s*Ald-Tx). Blue indicates functional transporters present in the three strains. Enzymes and transporters that are absent or nonfunctional in at least one of the three strains are indicated in red (exceptions indicated by superscript). Grey font indicates molecules that are not expected given the absence of the enzymes/transporters. Compare to *S. poulsonii* from *D. melanogaster*[121]. Created with BioRender.com.

**Supporting Figure S11.** Maximum Likelihood tree of the RIP (Ribosome Inactivating Protein) Domain sequences obtained with IQtree. Clade support values are shown at each node. Tip Labels are color coded by *Spiroplasma* clade. Bold-face tip labels are RIP genes found in the *Drosophila*-associated Citri clade strains *s*Ald-Tx, *s*Hy2, *s*Moj. MSRO = *s*Mel. Host species and major group (e.g. insect order) is also indicated.

**Supporting Figure S12**. Maximum Likelihood tree for each of the five blocks of the DNA alignment of members of Clade RAO. Tip Labels are color coded by *Spiroplasma* clade. Data from <u>Supporting Dataset S15</u>.

**Supporting Figure S13.** Maximum Likelihood tree of the amino acid alignment of the OTU domain of 81 *Spiroplasma* genes. Yellow highlighted clade is exclusive to *Drosophila*-associated *Spiroplasma* strains. Spaid = *s*Mel male killing gene[47]. Sequences were aligned with MAFFT and sites with at least 50% gapped taxa were removed. Input and output files are in <u>Supporting Dataset S16</u>.

**Supporting Figure S14.** Maximum Likelihood tree of the amino acid alignment of the ETX domain of 130 *Spiroplasma* genes obtained with IQtree. Sequences were aligned with MAFFT and sites with at least 50% gapped taxa were removed. Tip Labels are color coded by *Spiroplasma* clade. Bold-face tip labels are RIP genes found in the *Drosophila*-associated Citri clade strains *s*Ald-Tx, *s*Hy2, *s*Moj, and the Poulsonii clade *s*Hy1. Input and output files are in <u>Supporting Dataset S17</u>

**Supporting Figure S15**. Nucleotide alignment of the MTX-ETX2-domain containing genes of *s*Hy1 (00057), *s*Hy2 (EHV01_03630), and *s*Ald (EHU54_02490) and neighboring regions with high homology among two or all of these strains. Grey block arrow was found in a separate *s*Ald contig. CDS are marked with yellow block arrows. Citri clade strain name in red, whereas Poulsonii clade strain name in blue. Grey = identical to consensus; black = different from consensus. Alignment files are in <u>Supporting Dataset S18</u>.

**Supporting Figure S16**. Amino acid alignment of the PQQ domain region of 37 *Spiroplasma* genes. Residues are colored by amino acid, except for grey residues, which are not different from the consensus sequence. Sequence labels are colored by *Spiroplasma* clade: red = Citri; blue = Poulsonii; Grey = clade X. Genes containing also an ETX domain include the label “PQQ_ETX”. Alignment is Geneious format in <u>Supporting Dataset S19</u>.

**Supporting Figure S17.** Nucleotide alignment of *Spiroplasma Citri* clade regions with homology to lethal factor (or ADP-ribosyltransferase) and/or protective antigen (PA). Grey = identical to consensus; Black = different from consensus. Annotated coding sequences (CDS) are indicated by yellow block arrows below each sequence. Stop codons (*) are shown for some CDS. Conserved motifs or domains identified in coding sequences are indicated in magenta triangles or block arrows (R, STS, and QXE for ADP/lethal factor; and 3 domains for protective antigen). Phylogeny of Citri Clade (from Figure 2). Green in identity graph = identical nucleotide in all sequences covering that alignment position. Alignment in Geneious format in <u>Supporting Dataset S20</u>.

**Supporting Figure S18.** Maximum Likelihood tree of spiralin DNA sequences, retrieved from NCBI via blastn (or via “spiralin” term search for RefSeq annotated genomes at https://www.ncbi.nlm.nih.gov/datasets/gene), from members of the Citri (red) and Poulsonii (blue) clades. Grey: clade undetermined. Tree arbitrarily rooted with the spiralin C clade. Node support: SH-aLRT support (%) / aBayes support / ultrafast bootstrap support (%). Sequences were aligned with MAFFT (Algorithm = auto). Sites containing at least 80% gaps were removed. Files used to generate this tree are found in <u>Supporting Dataset S21</u>.

## Supporting Datasets Descriptions (all Supporting Datasets available at 10.6084/m9.figshare.c.7437997)

**Supporting Dataset S1**. Raw data obtained for the *Drosophila mojavensis Spiroplasma* protection experiments.

**Supporting Dataset S2**. Alignment of 16S rDNA sequences used for Supporting Figure S2. Three formats are included: fasta, phylip, and geneious.

**Supporting Dataset S3**. Excel Workbook containing several spreadsheets used to list tips in phylogenomics and phylogenetic trees. The spreadsheets contain the information used to replace tip labels in trees within FigTree.

**Supporting Dataset S4**. Alignment of 16S rDNA dataset used for Supporting Figure S2. Tree presented in Supporting Figure S2 is also included in Newick format.

**Supporting Dataset S5**. Input and output files of the Phylogenomics Analyses, including Orthofinder, Phipack, Mafft, and IQTree.

**Supporting Dataset S6**. Output files from cctyper.

**Supporting Dataset S7**. Pipeline and files used to search for genes annotated as plectroviruses.

**Supporting Dataset S8.** Alignment file (Geneious format) used to generate <u>Supporting</u> <u>Figure S5</u> comparing the phage-like contig of *Spiroplasma s*Hy2 (Citri clade) with those of the Poulsonii clade strains *s*Mel and *s*Neb reported in Ramirez et al. (2023).

**Supporting Dataset S9**. Output files for EggNog analyses of the *Drosophila*-associated Poulsonii clade strains *s*Mel, *s*Hy1, *s*Neo.

**Supporting Dataset S10**. KEGG IDs for the *Drosophila*-associated Citri clade strains *s*Ald-Tx, *s*Moj, and *s*Hy2.

**Supporting Dataset S11.** Data used to generate <u>Supporting Figure S8</u>. KEGG IDs and Brite Categories data for the *Drosophila*-associated Citri clade strains sAld-Tx, sMoj, and sHy2.

**Supporting Dataset S12**. KEGG Orthology numbers for the *Drosophila*-associated Poulsonii Clade strains *s*Mel, *s*Neo, and *s*Hy1, used for <u>Supporting Figure S9</u>.

**Supporting Dataset S13**. List of *Spiroplasma* strains encoding Trehalose A and Trehalose B genes.

**Supporting Dataset S14**. Files used for inferring the phylogeny of the RIP domain amino acid alignment shown in <u>Supporting Figure S11</u>. Contains alignment prior to masking and removing sequences that were too short, and IQTree input and output files.

**Supporting Dataset S15**. Files used to infer the phylogenies of the DNA alignment blocks of members of Clade RAO shown in <u>Supporting Figure S12</u>.

**Supporting Dataset S16**. Files used to infer the phylogeny of the OTU Domain amino acid alignment shown in <u>Supporting Figure S13</u>. Contains alignment before and after masking, and IQTree input and output files.

**Supporting Dataset S17.** Files used to infer the phylogeny of the ETX Domain amino acid alignment shown in <u>Supporting Figure S14</u>. Contains alignment before and after masking, and IQTree input and output files.

**Supporting Dataset S18**. Nucleotide alignment (Geneious format) of MTX genes used to generate <u>Supporting Figure S15</u>.

**Supporting Dataset S19**. Amino acid alignment (Geneious format) of the PQQ Domain used to generate <u>Supporting Figure S16</u>.

**Supporting Dataset S20.** Nucleotide alignment (Geneious format) of the genes annotated as lethal factor (or ADP-rybosyltransferase) and protective antigen used to generate <u>Supporting Figure S17</u>.

**Supporting Dataset S21.** Files used to infer the phylogeny of the Spiralin genes shown in <u>Supporting Figure S18</u>. Contains alignment before and after masking, and IQTree input and output files.

**Supporting Dataset S22.** Files used to identify genomes encoding (pseudo)gene for “1-acyl-sn-glycerol-3-phosphate acyltransferase” (Clades: Mirum, Chrysopicola, Citri, and Poulsonii). Files used to identify genomes encoding (pseudo)gene for “lysophosphatase” of “alpha/beta hydrolase” homologues (Clades: Citri and Poulsoni).

**Supporting Dataset S23.** Input and output files of Busco completeness analyses presented in Table 1.

## Supporting Protocols

**Supporting Protocol S1**. DNA Isolation using CTAB.

**Supporting Procotol S2**. DNA Isolation using Chloroform-Ethanol.

**Supporting Procotol S3.** Protocol to prepare *Drosophila* Opuntia-banana food media.

