## Supporting Table S5 for "Diverse toxin repertoire but limited metabolic capacities inferred from the draft genome assemblies of three *Spiroplasma* (Citri clade) strains associated with *Drosophila*"

**Supporting Table S5. *Spiroplasma* frequencies in wild-caught *D. aldrichi* and *D. mulleri* from sampled localities of Texas.**

| **Locality (Texas, USA)** | **Latitude** | **Longitude** | **Year** | **Species** | ***Spiroplasma*-positive** | ***Spiroplasma*-negative** | **Total** | **Frequency** |
| --- | --- | --- | --- | --- | --- | --- | --- | --- |
| Brackenridge Field Lab, Austin | 30.284751 | -97.778115 | 2013 | *D. aldrichi* | 31 | 2 | 33 | 0.94 |
| Brackenridge Field Lab, Austin | 30.284751 | -97.778115 | 2013 | *D. mulleri* | 24 | 8 | 32 | 0.75 |
| Freeman Ranch, San Marcos | 29.935278 | -98.014284 | 2012 | *D. aldrichi* | 13 | 10 | 23 | 0.57 |
| Freeman Ranch, San Marcos | 29.935278 | -98.014284 | 2012 | *D. mulleri* | 7 | 13 | 20 | 0.35 |
| Freeman Ranch, San Marcos | 29.935278 | -98.014284 | 2016 | *D. aldrichi* | 14 | 7 | 21 | 0.67 |
| Freeman Ranch, San Marcos | 29.935278 | -98.014284 | 2016 | *D. mulleri* | 1 | 25 | 26 | 0.04 |
| San Antonio | 29.676634 | -98.475748 | 2012 | *D. aldrichi* | 1 | 5 | 6 | 0.17 |
| San Antonio | 29.676634 | -98.475748 | 2012 | *D. mulleri* | 1 | 8 | 9 | 0.11 |
| **Total for *D. aldrichi*** | **all** | **all** | **all** | ***D. aldrichi*** | **59** | **24** | **83** | **0.71** |
| **Total for *D. mulleri*** | **all** | **all** | **all** | ***D. mulleri*** | **33** | **54** | **87** | **0.38** |
