## Supporting Table S6 for "Diverse toxin repertoire but limited metabolic capacities inferred from the draft genome assemblies of three *Spiroplasma* (Citri clade) strains associated with *Drosophila*"

**Supporting Table S6. P-values and assumed distribution (binomial or quasibinomial) from Analysis of Deviance Table (Type II tests) for the experiments that tested whether *Spiroplasma* *s*Moj influenced the interaction between *Drosophila mojavensis* and two species of parasitic wasps. None of the p-values were significant after Bonferroni correction for multiple comparisons (adjusted alfa = 0.05/11 = 0.0045).**

| **Wasp treatment** | **larva to adult fly survival** | | **larva to pupa survival** | | **wasp success**  **(adult wasp over fly larvae)** | | **pupal failure**  **(failed pupae over pupae)** | |
| --- | --- | --- | --- | --- | --- | --- | --- | --- |
| no wasp | 0.9554 | quasibinomial | 0.3434 | quasibinomial | n/a | n/a | 0.5153 | quasibinomial |
| Leptopilina_Lh14 | 0.9949 | binomial | 0.1191 | quasibinomial | 0.5552 | quasibinomial | 0.7988 | quasibinomial |
| Asobara_w35 | 0.1199 | quasibinomial | 0.04018 | quasibinomial | 0.504 | quasibinomial | 0.7941 | quasibinomial |
