## Supporting Table S2 for "Diverse toxin repertoire but limited metabolic capacities inferred from the draft genome assemblies of three *Spiroplasma* (Citri clade) strains associated with *Drosophila*"

**Table S2. Restriction enzymes and conditions to differentiate *Spiroplasma* strains *s*Hy1 (*Poulsonii* clade) and *s*Hy2 (*Citri* clade) at the 16S rRNA gene.**

| **Restriction Enzyme** | **Cut site** | **Temp (°C)** | ***s*Hy1 fragment sizes (bp)** | ***s*Hy2 fragment sizes (bp)** |
| --- | --- | --- | --- | --- |
| *Bsa*I | 5’...GGT CTC (1/5)^...3’ | 37 | 1,216  161 | No cut |
| *Sac*I | 5’...GAG CT/C...3’ | 37 | No cut | 818  550 |
