## Supporting Table S1 for "Diverse toxin repertoire but limited metabolic capacities inferred from the draft genome assemblies of three *Spiroplasma* (Citri clade) strains associated with *Drosophila*"

**Table S1. Insect species and strains/isofemale lines used for genome assembly and/or fitness assays.**

| **Insect Species** | **Strain or isofemale line (year established)** | ***Spiroplasma* strain** | **Location captured** |
| --- | --- | --- | --- |
| *Drosophila aldrichi* | Fr0317-09 (2017) | *Spiroplasma* sAld-Tx | San Marcos, Texas |
| *Drosophila hydei* | H25 (2014–2015) | *Spiroplasma* *s*Hy2 | Mexico |
| *Drosophila mojavensis* | CI-33-15 (2012) | *Spiroplasma* *s*Moj | Catalina Island, California |
| *Leptopilina heterotoma* | Lh14 (2002) | none | Winters, California |
| *Asobara sp.* | w35 (2012) | none | San Marcos, Texas |
