## Supporting Protocol S3 for "Diverse toxin repertoire but limited metabolic capacities inferred from the draft genome assemblies of three *Spiroplasma* (Citri clade) strains associated with *Drosophila*"

**Preparing fly food (Opuntia Banana recipe for 1 Liter)**

1L	deionized water

10g	agar

27.5g	yeast flakes

2g	Tegosept (same as Benzoic acid, p-hydroxy-, methyl ester; methyl paraben; nipagen; Lexgard M)

47.5g	Corn Syrup (we use HEB’s Hill Country Light Syrup)

30g	Malt

137.5g	bananas (not too ripe; we usually peel them and freeze them)

2.125g	Opuntia powder (We are currently using “Starwest Botanicals Nopal Cactus Powder Wildcrafted, 1 Pound”. We used to purchase fresh “nopalito” in cubes at the grocery store, dehydrate them in an incubator @ ~65°C, then grind them in a coffee grinder).

1.5g	Propionic Acid (add ~1ml of ethanol to dissolve; you can do it in a petri dish). The propionic acid is in a small tub under the sink. You can use the ethanol from the squirt bottle (usually in fly room).The Propionic acid plus ethanol mix is added once the food is cooling down (see below).

If (glass) vials have not been autoclaved, set the autoclave cycle **before** you begin preparing the food so that they will be ready.

Boil water for ~10 min in covered pot (you can re-use the aluminum foil to cover it)

Weigh all ingredients on Petri dishes except for corn syrup (use small beaker)

Place bananas, opuntia powder and malt in blender. With ladle take a few spoonfuls of the boiled water from the pot and add to blender. Blend and set aside.

Add agar, yeast flakes, corn syrup and Tegosept to pot with boiling water. Boil and mix with the whisk often for ~10 min. (use this time to prepare vials, to start cleaning/washing old vials, and/or to wash utensils)

Carefully pour the pot contents into the blender and blend at low speed (if you do high speed it will spill). Pour blender contents back into pot and boil (at low/med heat) for another ~10 min, mixing often.

Remove the pot from the heat and insert the food thermometer. In the meantime, add water the to the big pot and heat it up. It will be used as a water bath to keep the contents of the small pot warm while you pour vials.

Arrange the vials on clear racks or directly on the bench for pouring. They need to be visible so that you will put the correct amount.

Once the food has reached 145°F (63°C), mix in the Propionic Acid (plus ethanol).

Dispense the food into vials. We currently use a confectionery funnel (“Artilife Confectionery Funnel Stainless Steel with stand and three nozzles” to quickly dispense relatively similar amounts per vial efficiently. Othewise, you can use a ladle to serve food into a small beaker and from the small beaker pour into each vial. Refill the beaker as necessary.

Once all the food from the pot has been poured into vials, use the hot water from the large pot to begin cleaning utensils. Do not leave dirty dishes or benches.

Place the vials in the white plastic rack and cover with clean cheesecloth to prevent contamination by flies and minimize condensation. As soon as the food is settled, vials can be used. Once the vials are cool and their walls as are relatively free of condensation, cover with clean flugs, place the whole tray in a plastic bag (we reuse a white trash bag) and store in a refrigerator.
