## Supporting Protocol S1 for "Diverse toxin repertoire but limited metabolic capacities inferred from the draft genome assemblies of three *Spiroplasma* (Citri clade) strains associated with *Drosophila*"

CTAB Extraction Protocol

1. Place CTAB buffer in 60°C water bath for 30 min before use
2. Place tissue or specimen in 1.5-ml microtube
3. Add 300ul CTAB buffer to each tube
4. Grind specimen with pestle.
5. Cover tubes and incubate at 60°C for at least 1h
6. Add 300ul of sterile molecular grade water
7. Add 300ul of phenol, mix well, centrifuge for 10 min. Extract supernatant (top layer) and place in a new tube.
8. Add 300ul phenol-chloroform-isoamyl, mix well, and centrifuge for 10 min. Extract supernatant (top layer) and place in a new tube.
9. Add 300ul chloroform-isoamyl (24:1) to each tube, and shake by inverting for 2 min. Centrifuge for 10 min.
10. Label a new set of 1.5ml microtubes. Add 25ul 3M NaOAc and 600ul of 100% Ethanol (kept at -80C freezer) to each tube
11. Take aqueous (top) phase from step 9 and add to the tubes from step 10. Be sure not to disturb the interphase to avoid picking up lipids.
12. Place at -20°C for 20 min to overnight to precipitate
13. Centrifuge at high speed for 3 min
14. Decant fluid from DNA pellet and draw off remainder by pipetting (watch that the pellet does not float free from the bottom of the tube).
15. Add 200ul of 70% Ethanol and wash the pellet by gently inverting the closed tube several times, even if you have to flick the bottom of the tube to get the pellet to float free.
16. Centrifuge at medium to high speed for 2 minutes
17. Pour off ethanol and remove excess with pipette (do not dump pellet into waste)
18. Dry in oven at 65-70°C for ~5 min to remove excess Ethanol
19. Add 75ul of sterile water, 1x TE, or a low TE buffer such as Qiagen’s EB buffer.
20. Incubate at 65-70°C for 1 h to overnight to resuspend DNA.

**Reagents/consumables**

DWK Life Sciences Kimble™ Kontes™ Pellet Pestle™

Use to resuspend protein and DNA pellets or grind soft tissue in microcentrifuge tubes

Supplier:  DWK Life Sciences 7495200000

**CTAB Isolation buffer 2X**

Based on Coffroth et al. 1992. Marine Biology 114: 317-325

End concentrations: For 100 ml buffer mix:

1.4 M	NaCl	8.182g

20 mM	EDTA (pH 8)	0.74g

100 mM	Tris/HCl (pH 8)	1.21g

2% (w/v)	Hexadecyltrimethylammonium bromide CTAB powder	2 g

Add after filter sterilization under a hood:

0.2% (v/v)	β-mercaptoethanol (TOXIC!!!)	200 µl

- Add ddH_2_O to just under 100 ml.
- Warm to 65°C under stirring to bring the CTAB into solution.
- Once dissolved, bring final volume to 100 ml using a graduated cylinder.
- Filter sterilize (0.2 µm) into sterile 50 ml Falcon tubes and store at −20°C or 4C.
