## Supporting Protocol S2 for "Diverse toxin repertoire but limited metabolic capacities inferred from the draft genome assemblies of three *Spiroplasma* (Citri clade) strains associated with *Drosophila*"

**Chloroform-Ethanol DNA Purification Protocol**

**Reagents:**

Extraction buffer (described by volume for 1 sample)

Distilled water 0.85 mL

0.5 M EDTA 0.1 mL

10% SDS (Sodium dodecyl sulfate, vendor VWR) 0.05 mL

Potassium acetate 3M pH 4.2 (adjust pH with Acetic Acid glacial)

Chloroform

Isopropanol 100%

Ethanol 70%

**Procedure**

1. Add 1 mL of extraction buffer to each sample, homogenize and mix well

2. Put in 72$^{o}$ C water bath for 12 minutes, vortex and put in another 12 minutes

   Add 2 ul of RNase and incubate 5 minutes.

   Spin for 1 min (max speed)

3. Transfer supernatant to a new microcentrifuge tube with 50 µl of potassium acetate and mix well

4. Incubate on ice for 12 minutes, vortex and incubate another 12 minutes on ice

5. Spin 12 minutes at top speed (>13000 rpm)

6. Transfer supernatant into microcentrifuge tube with 500 µl of chloroform

7. Vortex well

8. Spin 3 minutes at top speed

9. Transfer 750 µl of the top phase into a new eppendorf with 500 µl of 100% IsoOH (mixing by inverting six times)

10. Spin 6 minutes at top speed

11. Discard supernatant

12. Add 1 mL of 70% EtOH

13. Spin 6 minutes at top speed

14. Discard supernatant

15. Tap dry on paper towel

16. Dry in Speed Vac for 12 minutes

17. Dissolve DNA in 100 µl TE buffer
